# Genome-wide association analysis of age-at-onset traits using Cox mixed-effects models

**DOI:** 10.1101/729285

**Authors:** Liang He, Alexander M. Kulminski

**Author notes:** Corresponding authors: Liang He, Alexander Kulminski.

## Abstract

Age-at-onset is one of the critical phenotypes in cohort studies of age-related diseases. Large-scale genome-wide association studies (GWAS) of age-at-onset can provide more insights into genetic effects on disease progression, and transitions between different stages. Moreover, proportional hazards or Cox regression generally achieves higher statistical power in a cohort study than a binary trait using logistic regression. Although mixed-effects models are widely used in GWAS to correct for population stratification and family structure, application of Cox mixed-effects models (CMEMs) to large-scale GWAS are so far hindered by intractable computational intensity. In this work, we propose COXMEG, an efficient R package for conducting GWAS of age-at-onset using CMEMs. COXMEG introduces fast estimation algorithms for general sparse relatedness matrices including but not limited to block-diagonal pedigree-based matrices. COXMEG also introduces a fast and powerful score test for fully dense relatedness matrices, accounting for both population stratification and family structure. In addition, COXMEG handles positive semidefinite relatedness matrices, which are common in twin and family studies. Our simulation studies suggest that COXMEG, depending on the structure of the relatedness matrix, is 100∼100,000-fold computationally more efficient for GWAS than coxme for a sample consisting of 1000-10,000 individuals. We found that using sparse approximation of relatedness matrices yielded highly comparable performance in controlling false positives and statistical power for an ethnically homogeneous family-based sample. When applying COXMEG to a NIA-LOADFS sample with 3456 Caucasians, we identified the *APOE4* variant with strong statistical power (p=1e-101), far more significant than previous studies using a transformed variable and a marginal Cox model. When investigating a multi-ethnic NIA-LOADFS sample including 3456 Caucasians and 287 African Americans, we identified a novel SNP rs36051450 (p=2e-9) near *GRAMD1B*, the minor allele of which significantly reduced the hazards of AD in both genders. Our results demonstrated that COXMEG greatly facilitates the application of CMEMs in GWAS of age-at-onset phenotypes.

## Introduction

Age-at-onset and time-to-event are among the most important phenotypes of interest in cohort studies of age-related diseases (He et al., 2016a; Kulminski et al., 2016), and behavioral genetics (He et al., 2016b) as these phenotypes carry information on timing of events. Genome-wide association studies (GWAS) of time-to-event phenotypes are critical for gaining insights into the genetics of disease development and progression. Specifically, for many age-related diseases with unknown causal mechanisms, such as Alzheimer’s disease (AD), pathological processes may be triggered decades before the first symptoms. Identifying genetic variants associated with age-at-onset may provide insights into potential etiology of such diseases. Time-to-event phenotypes are also of inherent interest in specific fields such as clinical trials. In such situations, time-to-event analysis is essential because events occur in a subset of subjects whereas information on the outcome for other subjects is mostly censored. Moreover, previous evidence shows that cohort studies using proportional hazards (or Cox) regression models generally increase statistical power compared to case-control studies using logistic regression model (Callas et al., 1998; Green and Symons, 1983; Staley et al., 2017). The latter may also yield inflated estimates of the effect sizes in GWAS (Staley et al., 2017).

While time-to-event analysis using a Cox model plays a central role in many biological and epidemiological studies and is often more powerful, its application to GWAS is hampered by its computational burden when random effects accounting for unexplained heterogeneity (e.g., genetic relationship) are included. Unlike a computationally efficient linear mixed model (LMM), which has been well optimized and broadly applied in genetics (Kang et al., 2008; Lippert et al., 2011; Loh et al., 2015; Svishcheva et al., 2012; Zhou and Stephens, 2012), much less attention has been paid to Cox mixed-effects models (CMEMs). To facilitate genetic association analysis in family-based studies, (Therneau, 2015) developed a specific data structure (bdsmatrix) and a fast algorithm implemented in C in the Cox mixed-effects R package (coxme), optimized for block-diagonal kinship matrices, which are constructed based on pedigree information (Pankratz et al., 2005; Therneau, 2003). Our analyses show that the performance of the coxme package is remarkable for block-diagonal kinship matrices, reducing the CPU time for one single nucleotide polymorphisms (SNP) from ∼15 minutes for a dense relatedness matrix to less than one second when analyzing a medium cohort having ∼3500 individuals (e.g., Late-Onset Alzheimer’s Disease Family Study from the National Institute on Aging (NIA-LOADFS) (Lee et al., 2008)). Despite this dramatic improvement, applying a CMEM (Ripatti and Palmgren, 2000; Therneau, 2015) to a large-scale GWAS involving millions of SNPs is still computationally challenging. Even for the NIA-LOADFS, it would take coxme around three months using one CPU thread to run a GWAS for 10M imputed SNPs. Moreover, it is limited to block-diagonal and positive definite relatedness matrices. For a more general genetic relationship matrix (GRM) (e.g., the GRM proposed in (Yang et al., 2011)) that cannot be grouped into a block-diagonal matrix or is not positive definite due to e.g., monozygotic (MZ) twins, coxme would take decades or fail.

Current GWAS often include a large number of subjects and studies, especially in consortia, and interrogate higher-order relationships such as SNP-environment and SNP-SNP interactions, which requires a large number of tests. Therefore, there is an urgent need to develop fast algorithms for a CMEM that can be applied to large-scale GWAS involving time-to-event traits. In this work, we introduce efficient estimation and testing algorithms for a CMEM through an R package COXMEG (https://r-forge.r-project.org/R/?group_id=2366), in which G stands for GWAS. COXMEG addressed the aforementioned computational problems, and is well suited for large-scale GWAS. Our model has five major improvements. First, it substantially improved computational efficiency. For example, for a family-based design using a block-diagonal kinship matrix, COXMEG is often ∼100-fold faster than for the remarkably optimized coxme package in the same computational environment. Second, we developed an inexact Newton method in COXMEG, which is not restricted to a block-diagonal pedigree-based relatedness matrix. In general, a sparse matrix or any matrix that can be approximated by sparsity (e.g., an autoregressive (AR) matrix using banding) is able to enjoy the computational benefit from COXMEG, which significantly expands the application of COXMEG. For example, we showed in our application to NIA-LOADFS that COXMEG attains high computational performance and statistical power using a sparse GRM after thresholding. Third, when a fully dense relatedness matrix cannot be approximated by a sparse matrix, it is computationally demanding to estimate the hazard ratios (HRs) and then perform a Wald test to obtain p-values. For such situations, we propose a fast score test, which obtains the p-values without estimating the effect size of each SNP. Fourth, COXMEG generalizes the CMEM to handle positive semidefinite relatedness matrices, which are very common in GWAS. Fifth, for a large-scale (>20,000) fully dense relatedness matrix, we introduce a fast randomized algorithm in COXMEG to estimate the variance component, which considerably reduces computational time.

Feasibility of COXMEG was tested by performing GWAS of age-at-onset of AD in NIA-LOADFS. AD is a complex age-related disease, which spans multiple stages and probably involves different biological mechanisms (e.g., beta-amyloid deposition, pathogenesis, vascular changes, activation of the immune system and phosphorylation of tau (Finch & Kulminski, 2019)) in different stages. Our hypothesis is that genetic architecture of AD is likely implicated in faster pathological progression from the initial stage. We demonstrate high computational performance and report novel findings using a trans-ethnic sample of NIA-LOADFS.

## Results

### Overview of COXMEG

Depending on whether the relatedness matrix is sparse, we propose in COXMEG two algorithms accounting for the dependence in subjects, COXMEG-sparse for a sparse relatedness matrix, and COXMEG-score for a dense relatedness matrix. Two primary purposes to adjust for dependence structure in the context of GWAS are (i) correcting for population stratification, (ii) accounting for family structure. We consider these two purposes separately because the rationale and thus the relatedness matrices used in the two situations are different. In the first case, the major concern is to avoid identifying false positive SNPs correlated with race. This is because population stratification results in many SNPs that can almost tag the race information in the sample, and consequently are highly correlated with race. If race or any relevant variables, such as principal components (PCs), is not included in the model, the estimated effects of these SNPs should be attributed to the effect of race, leading to a large number of false positive SNPs. In the second case, however, the major benefit of explicitly specifying the sample dependence is to control the overall false positive rate and increase statistical power due to family structure. Previous studies show that ignoring dependence in the Cox model leads to overestimate of the model parameters (Wei et al., 1989). We show in our analyses that COXMEG-score is well suited for the first case, and COXMEG-sparse is dedicated to the second case.

Sparse approximation of the relatedness matrix is attractive. When a selected study population is almost ethnically homogeneous and a CMEM is adopted primarily to correct for family structure, for example, in family-based GWAS (He et al., 2016a), we found that the relatedness matrix is often sparse or can be approximated by a sparse matrix. For a family-based study, a widely-used relatedness matrix is a pedigree-based kinship matrix, which is generally block-diagonal and sparse. When a GCTA GRM is used (Yang et al., 2011), which is computed from genotypes, we noticed that it often has a large proportion of near-zero elements. We show in the following analyses that, compared to a fully dense GRM, it often produced close results to use a sparse approximation of the GRM by thresholding the whole matrix or extracting its block-diagonal entries. However, using a sparse approximation is computationally much less intensive in terms of both speed and memory usage. Compared to coxme, which is only optimized for a block-diagonal pedigree-based kinship matrix, COXMEG-sparse provides efficient algorithms for more general sparse matrices, which can also be positive semidefinite.

### COXMEG-sparse efficiently corrects for dependence in family-based study

To investigate the computational efficiency of COXMEG-sparse for block-diagonal relatedness matrices, we compared it with the coxme R package (Therneau, 2015), which is widely used to fit a CMEM and highly optimized for block-diagonal matrices. We also included in the comparison the function ‘coxph’ in the survival R package (Therneau and Lumley, 2015) with a shared Gaussian frailty model. A shared frailty model (Vaupel et al., 1979) was used as a fast but generally less accurate alternative to coxme in some studies (He et al., 2016b). It assumes that all members in one family share the same genetic component, which might not be a realistic presumption in many situations. In both coxme and coxph, block-diagonal approximation was used. Our simulation results (Figure 1) showed that it took COXMEG-sparse ∼60s (using one CPU thread) to analyze 10k SNPs for a small sample of 2000 individuals with a family size of 5. In contrast, under the same computational configuration, it took coxme ∼200min for the same task, the running time of which is ∼200-fold as much as that of COXMEG-sparse. We found that COXMEG-sparse was ∼20-fold faster than coxph under this setting. For block-diagonal relatedness matrices, the computational burden of all three methods rose almost linearly with increasing sample size (Figure 1), which was consistent with theoretical estimates given in the Methods section. When the sample size was up to 10,000, COXMEG-sparse was >1000-fold and >100-fold faster than coxme and coxph, respectively (Figure 1). We noticed that the computational burden of COXMEG-sparse increased more rapidly than coxme with respect to the family size, but it was still far below that of coxme when the family size was 100, which is rarely encountered in a real study (Figure 1). On the contrary, the computational burden of coxph dropped rapidly with increasing family sizes. This is because coxph depends on the number of families, which decreased when the total sample size kept unchanged.

**Figure 1:**
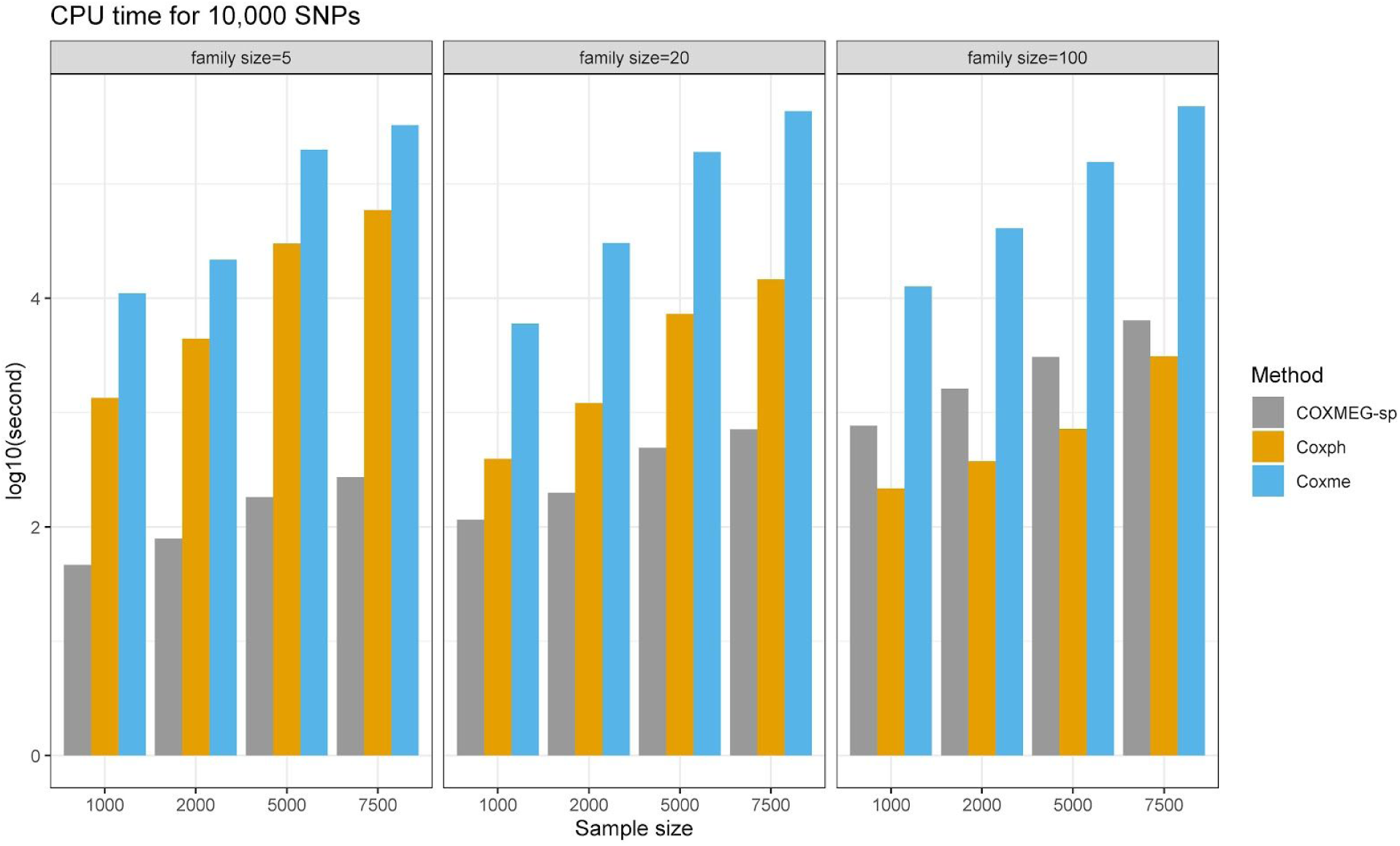
Comparison of computational time using one CPU thread (log_10_ (second)) of analyzing 10,000 SNPs using COXMEG-sparse, coxph and coxme under different sample sizes and family sizes with a block-diagonal relatedness matrix. Computational time of COXMEG-sparse includes estimation of the variance component and SNP effects.

### COXMEG-sparse is fast for general sparse matrices

Next, we examined the performance of COXMEG-sparse for more general sparse relatedness matrices. We assessed a class of sparse GRMs which were obtained by approximating a dense GRM by hard-thresholding small values so that individuals with long genetic distance were treated as independent. This type of GRMs is of our interest because we noticed that most elements in a GRM based on an infinitesimal model are almost zero when conducting a family-based study consisting of an ethnically homogeneous cohort. For instance, when we studied only the Caucasian subjects in the NIA-LOADFS, and set all elements below a prespecified small threshold (e.g., 0.02) to zero, the resulting sparse GRM had on average <1% non-zero elements per row, and most of them are from the family members. This type of GRMs is not block-diagonal because it has sporadic non-zero elements. We observed that COXMEG-sparse was dramatically more efficient to deal with this kind of approximated GRMs than coxme, which was extremely slow for a matrix that is not block-diagonal. By thresholding a GRM from the NIA-LOADFS using different cutoffs, we examined sparse GRMs, which had ∼3500 subjects and 20-30 non-zero elements per row. Our results show that it took coxme ∼17m for estimating only one SNP, while it took COXMEG-sparse only ∼0.01s when 1% of the elements were non-zero (Figure 2A). The computational time decreased with increasing sparsity of the relatedness matrix (Figure 2A).

**Figure 2.**
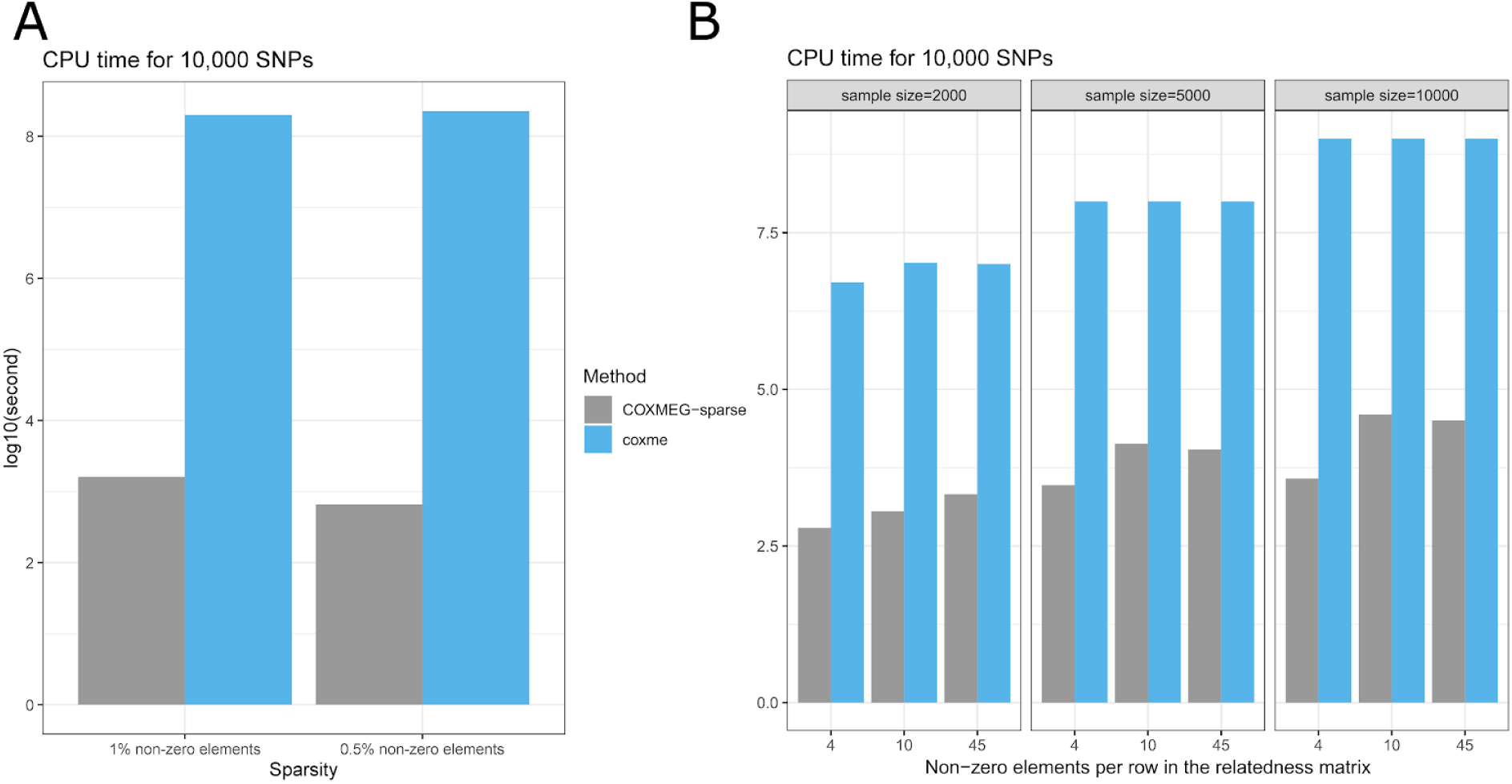
Comparison of computational time (log_10_(second)) of analyzing 10,000 SNPs using COXMEG-sparse and coxme under different sparsity patterns (0.1-2% non-zero elements) and/or sample sizes. Computational time of COXMEG-sparse includes estimation of the variance component and SNP effects. (A) CPU time for sparse relatedness matrices obtained by hard-thresholding a GRM constructed using the genotypes of 3456 Caucasian subjects from the NIA-LOADFS; (B) CPU time for sparse relatedness matrices obtained by banding an AR(1) correlation matrix. The computational time of coxme under 7500 and 10,000 samples were projected as it would take coxme too long to run one SNP.

To evaluate the performance of COXMEG-sparse for other correlation patterns, we conducted a simulation study using banded first-order autoregressive (AR(1)) matrices, which cannot be factorized as a block-diagonal matrix. Similarly, we found that COXMEG-sparse was substantially computationally faster than coxme (Figure 2B). The computational time increased almost linearly with respect to the sample size, much slower than coxme (Figure 2B). We did not include coxph in the above analyses because a shared frailty component is not clearly defined for a general sparse covariance matrix.

### Efficient association test using COXMEG-score for general relatedness matices

When there is a necessity to correct for both family structure and population stratification in a family-based multi-ethnic study, the GRM is usually dense and cannot be approximated by a sparse matrix using naive thresholding methods such as hard thresholding. For a fully dense GRM, the estimation algorithm would be extremely slow for coxme, which may even take 10 minutes to analyze one SNP with a small sample size of 2000. For such dense GRMs, we implemented a preconditioned conjugate gradient (PCG) method, termed COXMEG-pcg, which substantially reduces computational intensity compared to the Cholesky decomposition used in coxme. Our simulation studies indicated that its computational time dropped drastically compared to coxme in practice (Figure 3A). However, the efficiency of COXMEG-pcg might still not be acceptable when the speed is a leading concern of the application. We thus propose COXMEG-score, which is much faster than COXMEG-pcg by conducting a score test to obtain the p-values. We found in our simulation study that COXMEG-score completed the analysis for 10k SNPs of a sample consisting of 1000 individuals within 15s, computationally much more efficient than COXMEG-pcg and coxme, which took 10,00s and 10,000s, respectively (Figure 3A). When the sample size increased up to 10,000, it took COXMEG-score and COXMEG-pcg ∼1.4 hours and ∼1.2 days, respectively, while it would take coxme ∼3 years according to our projection. Nevertheless, it is unfortunately hard in this case to both efficiently obtain HRs and perform the test although HRs are often of interest in follow-up downstream analyses such as meta-analysis and causal inference.

**Figure 3.**
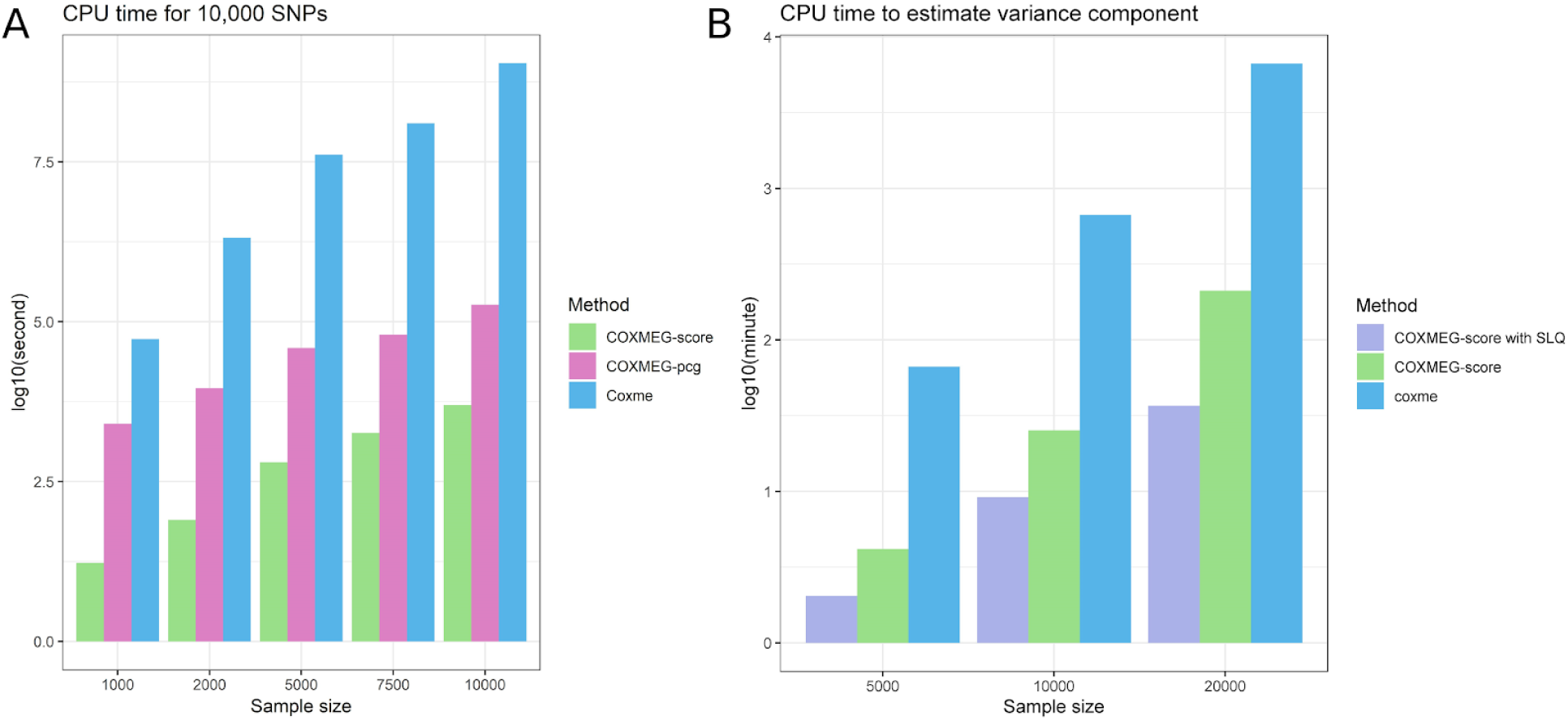
(A) Comparison of computational time (log_10_ (second)) of analyzing 10,000 SNPs using COXMEG-score, COXMEG-pcg, and coxme with a dense relatedness matrix of different sample sizes; (B) Comparison of computational time (log_10_ (minute)) of estimating the variance component using COXMEG-score with the exact log-determinant, COXMEG-score with a SLQ approximation, and coxme with a dense relatedness matrix of different sample sizes. The CPU time of coxme for a sample size >5,000 was projected.

For a very large cohort (e.g., >20,000), the estimation of the variance component in COXMEG-score becomes a computational bottleneck because the estimation requires an evaluation of a log-determinant of a large matrix, whose time complexity is cubic. For large cohorts, we introduce a randomized method using stochastic lanczos quadrature (SLQ) (Bai et al., 1996; Golub and Meurant, 2009; Ubaru et al., 2017) to approximate the log-determinant, which reduce the time complexity to quadratic. Our results show that COXMEG-score with a SLQ method employing 100 Monte Carlo samples and 10 steps of the Lanczos algorithm markedly lowered the computational burden for a cohort with >10,000 individuals (Figure 3B), and maintained the control of false positive rate (FPR) of detecting significant SNPs (Supplementary materials A.4). Our simulation results indicate that the accuracy of the SLQ approximation highly depended on the structure of the relatedness matrix, and an estimated variance component was particularly accurate for a large relatedness matrix with weak correlations (i.e., more accurate for a larger sample size and a smaller condition number of the relatedness matrix) (Supplementary materials A.4).

### COXMEG controls false positives and is more powerful to identify *APOE4* associated with AD

We first performed simulation studies to assess statistical power and FPR of COXMEG. We found that COXMEG, including COXMEG-score and COXMEG-sparse, had nearly the same statistical power and FPR as coxme, while coxph with a shared frailty had inflated FPR in certain situations of family-based datasets (Figures S1, S2). Detailed results about the evaluation of the statistical power and FPR can be found in Supplementary materials A.5.

Next, we performed GWAS of age-at-onset of AD for the NIA-LOADFS dataset to detect SNPs associated with the hazards of late-onset AD. We first focused on a homogeneous subpopulation by selecting 3456 Caucasian subjects, whose information about AD status was available, from the cohort. For the subjects whose information about the age-at-onset of AD was missing, we treated them as censored and set its age-at-onset as the age at the recruitment. The genotypes were imputed using Michigan Imputation Server with a reference panel from HRC (Version r1.1 2016) (Das et al., 2016). A total of ∼5.42M SNPs were interrogated in the analyses after removing SNPs having minor allele frequency (MAF)<0.05. Sex was included as a covariate in all analyses.

In addition to identifying variants associated with the onset of AD, we investigated the performance of COXMEG in the following three aspects. First, we evaluated the performance of COXMEG-sparse, COXMEG-score, coxme and coxph (with a shared Gaussian frailty corresponding to the families) in terms of statistical power and control of the genomic inflation factor. Second, we investigated potential influence on the statistical power of using different approximated GRMs in COXMEG-sparse. Third, we investigated the performance of using a uniform variance component estimated from the null model for analyzing all SNPs. More specifically, we assumed in coxph that all members in each family shared the same frailty. In the other three methods, we computed a GRM using all qualified genotypes (filtered by MAF >0.05 and missing rate <1e-4) from a 600k Illumina Human610_Quadv1_B genotype array. The GRM was estimated based on the GCTA model (Yang et al., 2011) (also see e.q. (10) in the Methods section) and was further scaled using the SNPRelate R package (Zheng et al., 2012) so that the diagonal elements equaled one. For coxme, we used a block-diagonal GRM by first defining diagonal blocks according to the families, and then setting all the elements outside the block-diagonal parts to zero. We found that eight pairs of Caucasian individuals were in complete correlation, indicating that they were MZ twins. As coxme does not handle such positive semidefinite matrices, we set those off-diagonal elements corresponding to the MZ twins to 0.99 to make the GRM positive definite. For COXMEG-sparse, we tested two GRMs, (i) the same block-diagonal GRM as used in coxme, and (ii) a sparse GRM by setting all elements in the GRM that had an absolute value <0.03 to zero. For COXMEG-score, we tested three GRMs, the original GRM, the block-diagonal GRM and the sparse GRM. In terms of the computational efficiency, the analyses using COXMEG-sparse and COXMEG-score were completed in ∼1 day using one CPU thread, while it took coxph and coxme more than three weeks to complete the analysis. Surprisingly, we found that coxph took longer than coxme.

Our results show that the *APOE4* variant was the strongest SNP, associated with increasing hazards of AD with a p-value of 1e-101 (Figure 4A), which was more significant than the p-value (p=3.3e-96) reported from a previous study of age-at-onset of AD using a much larger sample size (9162 Caucasian participants) (Naj et al., 2014), in which the NIA-LOADFS sample is a subset. The previous study used log-transformed age-at-onset as the phenotype and generalized estimating equations for estimation (Naj et al., 2014). This might suggests the CMEM is a more appropriate and powerful model than a marginal model in this case. In addition, we found that rs36051450, located on 11q24.1 and ∼25kb upstream from *GRAMD1B*, yielded a p-value of 8e-8, narrowly below the genome-wide significance (Figure 4A). *GRAMD1B* is a cholesterol transporter mediating non-vesicular transport of cholesterol from the plasma membrane to the endoplasmic reticulum. In contrast, a logistic mixed effects model produced a p-value of only 2.85e-05 for rs36051450. We also examined 23 common SNPs identified by a recent large-scale meta-analysis associated with incidence of AD (Jansen et al., 2019). We found that eight SNPs (in *CR1, BIN1, CLNK, HLA-DRB1, MS4A6A, PICALM, ABI3, CASS4*) were nominally significantly (p<0.05) associated with the hazards of AD (Table S1), in which rs4663105 in the *BIN1* region showed the strongest significance (p=2.961e-04).

**Figure 4.**
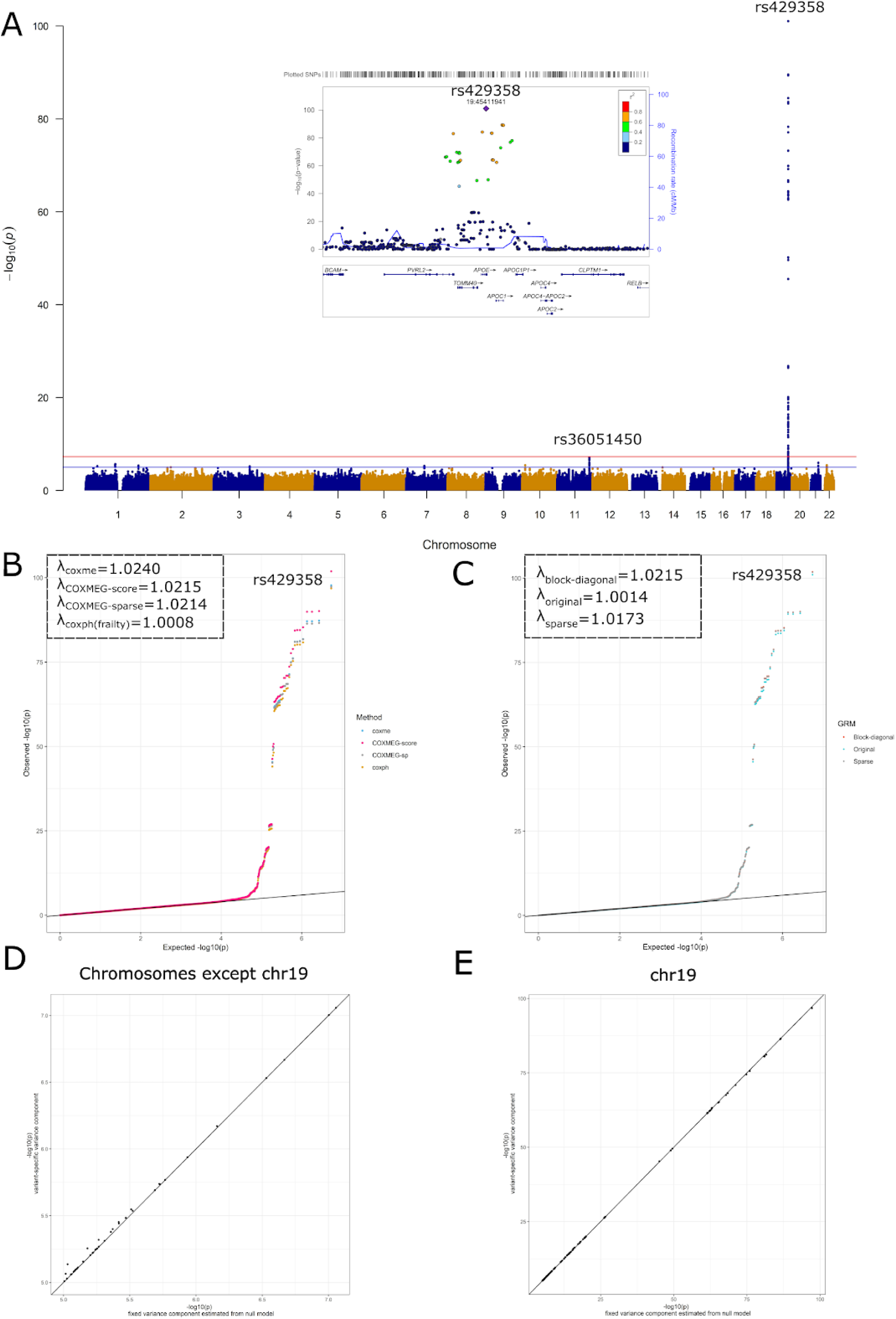
Results of GWAS of age-at-onset of AD with a LOADFS Caucasian sample. (A) P-values from GWAS of age-at-onset of AD using COXMEG-score with a GCTA GRM estimated from a genotype microarray. The local plot shows the p-values of significant SNPs in the *APOE* region. The red horizontal line is the genome-wide significance (5e-8) and the blue horizontal line is the suggestive significance (1e-5). (B) Comparison of the p-values from GWAS of age-at-onset of AD with a LOADFS Caucasian sample using four different methods (coxph with a shared frailty, coxme, COXMEG-sparse, COXMEG-score). A block-diagonal GRM reflecting family structure was used for the latter three methods. Genomic inflation factors are shown on the top left corner. (C) Comparison of the p-values from GWAS of age-at-onset of AD with a LOADFS Caucasian sample using three different GRMs (a raw GCTA GRM (original), a block-diagonal matrix obtained by extracting the block-diagonal part of the GCTA GRM reflecting family structure, a sparse matrix by thresholding the GCTA GRM with 0.03). Genomic inflation factors are shown on the top left corner. (D), (E) Comparison of p-values using a fixed variance component estimated from the null model and a variant-specific variance component for SNPs with p<1e-5 outside chromosome 19 (D) and on chromosome 19 (E).

Our results further showed that COXMEG-sparse, COXMEG-score, coxme using the same block-diagonal GRM, and coxph had almost the same genomic inflation factor (λ=1.021368, 1.021491, 1.023954, and 1.00083 respectively) (Figure 4B), suggesting that all four methods were comparable in terms of controlling false positives for this analysis. Using a family-based block-diagonal GRM or assuming a shared frailty was not inflating the p-values in this study. We observed that coxph produced slightly less significant p-values for top SNPs than the other methods. On the other hand, we observed that the statistical power of COXMEG-score to detect the *APOE4* variant was much higher (>four order of magnitude) than that of the other three methods (Figure 4B). We observed little difference in the p-values for detecting top SNPs between the original GRM, the approximated sparse GRM and the block-diagonal GRM (Figure 4C). In addition, they yielded almost the same genomic inflation factor (λ=1.00141, 1.021491, and 1.01727, respectively), suggesting that the sparse approximations were practically viable for ethnically homogeneous family-based cohorts. Finally, we evaluated potential consequences of using a fixed variance component estimated from the null model instead of a variant-specific variance component for top SNPs. We estimated a variant-specific variance component, and recomputed the effect size and its p-value for each SNP having a p-value<1e-5. We found that using the fixed variance component did produce slightly less significant p-values for those SNPs outside the *APOE* region (Figure 4D), while no evident difference was observed for those SNPs in the *APOE* region (Figure 4E). However, the variance component estimated from the null model including only sex was 0.26, while it rose to 0.38 when the null model included both sex and the *APOE4* variant (rs429358).

### COXMEG identifies novel SNPs associated with age-at-onset of AD

We applied COXMEG-score to a sample consisting of both ∼3500 Caucasians and 287 African Americans in NIA-LOADFS. The genotypes were imputed using Michigan Imputation Server with the reference panel from 1000 Genomes Phase 3 (Version 5) (Das et al., 2016). A total of 20M SNPs were included in the GWAS of age-at-onset of AD after removing SNPs having MAF<0.05. The GRM was constructed using the same procedure as in the previous analyses. When applying COXMEG-score, we included sex as a covariate. To examine the consequence without adjusting for the dependence between subjects, we performed two additional GWAS using coxph without frailty. In the first analysis, we included only sex as a covariate. In the second analysis, we included sex and top five PCs computed from the GRM to account for population stratification. We did not include coxph with a shared frailty or coxme because they were too time consuming to finish the analysis in a reasonable time.

Given the censoring proportion is 55%, the heritability of the risk of AD based on the common SNPs was 0.43 using e.q. (8). The genomic inflation factors were 1.01357, 1.06339, and 1.06493 for COXMEG-score, coxph and coxph with PCs, respectively (Figure 5A), suggesting that the false positive rate (FPR) was inflated when not accounting for the family structure. Using PCs to account for population stratification slightly improved the FPR. We observed that COXMEG-score yielded more significant p-values for the top SNPs than coxph with PCs (Figure 5A), suggesting that COXMEG-score achieved higher statistical power to detect SNPs in the APOE region. The top SNP was again the *APOE4* SNP rs429358 with a p-value of 1e-91 (Figure 5B), slightly less significant than that obtained from the Caucasian sample alone, which probably indicates that the effect of *APOE4* is lower in African Americans than Caucasians. Outside the *APOE* region, we identified a novel SNP rs36051450 (p=2e-9), and the p-value was more significant than that from the Caucasians alone. We found that the minor allele of rs36051450, with an allele frequency of ∼18% in the sample, had a protective effect on AD, significantly reducing the hazards in both male and female individuals (Figure 5C). The effect in male was less significant than that in female probably because of the smaller sample size of male. Interestingly, rs735665, a nearby SNP in high LD (r^2^=0.94) with rs36051450, is significantly (p-value=4e-9 to 4e-39) associated with chronic lymphocytic leukemia (CLL) and follicular lymphoma (Berndt et al., 2013; Conde et al., 2010; Di Bernardo et al., 2008; Slager et al., 2012; Speedy et al., 2014). In contrast, the minor allele of rs735665 increases the risk of CLL (OR=1.45-1.81), which suggests a potential antagonistic pleiotropy of these SNPs.

**Figure 5.**
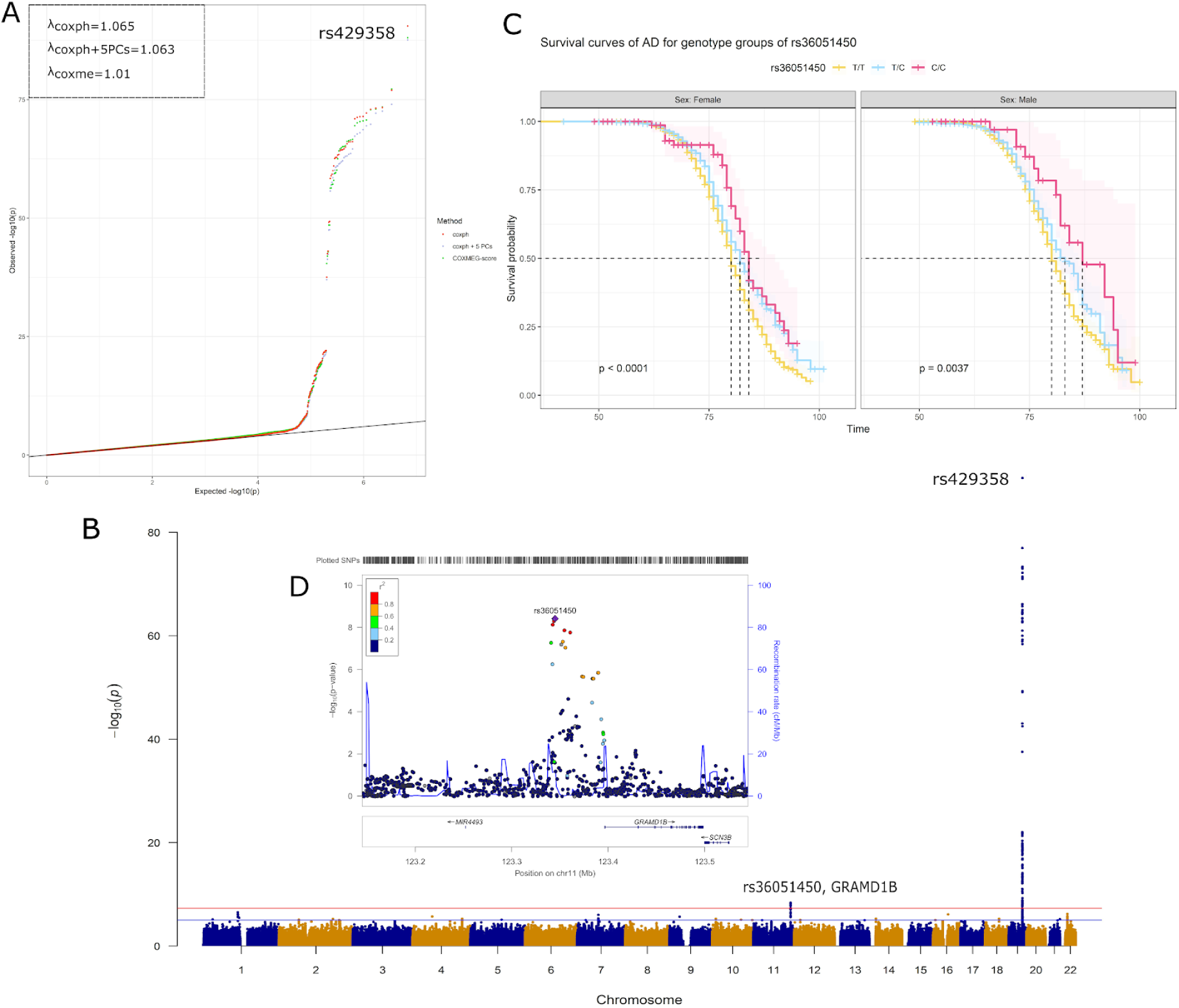
Results of GWAS of age-at-onset of AD with a sample consisting of Caucasians and African Americans in LOADFS. (A) Comparison of the p-values using three different methods (coxph, coxph with 5 top PCs estimated from a GCTA GRM as covariates, COXMEG-score with a GCTA GRM). Genomic inflation factors are shown on the top left corner. (B) P-values from GWAS of age-at-onset of AD using COXMEG-score with a GCTA GRM estimated from a genotype microarray. The red horizontal line is the genome-wide significance (5e-8) and the blue horizontal line is the suggestive significance (1e-5). (C) Survival curves of AD in three genotype groups of rs36051450 in men and women. (D) A plot shows the p-values of SNPs in the local region of rs36051450.

## Discussion

Age-at-onset is one of the common and fundamental traits in genetic epidemiology. When the underlying hazard function is unknown, the Cox model (Cox, 1972) is a prevalent routine for analyzing age-at-onset because it is more flexible than a parametric hazard model (e.g., Weibull). When subjects are correlated, a CMEM is frequently used for modelling the dependence. Although the CMEM has been investigated for decades (Cortiñas Abrahantes and Burzykowski, 2005; Ha et al., 2001; McGilchrist, 1993; McGilchrist and Aisbett, 1991; Ripatti and Palmgren, 2000; Ripatti et al., 2002), unfortunately, its estimation has no closed form in contrast to a parametric hazard frailty model, in which it is possible to express the marginal likelihood explicitly using the Laplace transform for some frailty families (Wienke et al., 2003; Yashin et al., 1995). Iterative methods are required for estimation, which are computationally intensive, and consequently prohibit its broad application in GWAS. (Therneau, 2003, 2015) optimized the estimation algorithm specifically for block-diagonal covariance matrices, and achieved remarkable performance, but its use is still limited in large-scale GWAS.

This work sets out to fill in the gap in the application of the CMEMs in GWAS. We have proposed efficient algorithms in COXMEG to facilitate the application of the CMEM to genome-wide age-at-onset or time-to-event analyses. We demonstrated in our simulation and real data analyses that COXMEG substantially improves computational efficiency, and is applicable for a wider range of relatedness matrices. Specifically, for a block-diagonal relatedness matrix, COXMEG is often hundreds to thousands of times faster than coxme, while for a fully dense relatedness matrix, COXMEG using a score test can be 100,000-fold faster. In addition, compared to coxme, COXMEG deals with positive semidefinite relatedness matrices, which are common in the context of GWAS. We demonstrated in a study of AD using a sample from the NIA-LOADFS that these improvements and generalizations in COXMEG allow fast GWAS of age-at-onset for a cohort of thousands or tens of thousands of individuals. For example, COXMEG completed the genome-wide analysis within one day for 10M SNPs in NIA-LOADFS (∼3500 subjects) using one CPU thread. Our work will be of particular interest to many areas in epidemiology such as age-related diseases (He et al., 2016b; Kulminski et al., 2016), behavior genetics (He et al., 2016a) and perhaps clinical trials, in which age-at-onset is a central concern. Moreover, the application of COXMEG includes, but is not limited to genetic data. As demonstrated, COXMEG-sparse is computationally fast for general sparse matrices. A time-to-event analysis involving modeling dependence structure, e.g., geographically clustered time-to-event data, can benefit from COXMEG.

Our results about the risk of AD show that the CMEM may identify new SNPs that are not significant using a linear or logistic model. Interestingly, for the identified novel SNP rs36051450 near *GRAMD1B*, the p-value from the logistic model was less significant than that from the CMEM, which suggests that this SNP might contribute more to accelerating the disease progression than the incidence. According to GTEx (GTEx Consortium et al., 2017), *GRAMD1B* is abundantly expressed in brain tissues. Cell-sorting experiments of human cortex tissue indicates that *GRAMD1B* is most strongly expressed in astrocytes and neurons among five major brain cell types (Zhang et al., 2016). Further research could be conducted to replicate the findings in other cohorts.

Analogous to heritability defined in a linear model, a generalized heritability defined on a logarithmic scale of cumulative hazards can be obtained as proposed in (Korsgaard et al., 1999; Schneider et al., 2005; Yazdi et al., 2002) using e.g., the following formula,

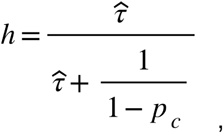

where *p*_*c*_ is the proportion of censored subjects and 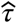 is the estimated variance component in CMEMs.

Despite the efforts made in COXMEG, it is still challenging in terms of both computational time and space to handle a fully dense relatedness matrix for a cohort of >50,000 individuals in a trans-ethnic study. As discussed in (Gianola et al., 2016), a fast and viable solution to account for sample dependence is to split the relatedness matrix into two parts based on PCA. As we showed in our analyses of AD using the NIA-LOADFS, using a sparse approximation of the GRM in an ethnically homogeneous study of family-based design achieved highly comparable performance. However, using a sparse approximation considerably alleviates computational burden. Thus, an efficient solution to a very large cohort might be to use top PCs from the GRM to correct for population stratification, and a sparse approximation to correct for family structure. In addition, we point out that that although COXMEG can handle a positive semidefinite kinship or GRM matrix, the number of zero eigenvalues has to be small. Otherwise, there would be a convergence problem because you would have too many free parameters to estimate without a penalty. Future work needs to be carried out to validate its application to twin studies.

## Methods

### Model specification in COXMEG

We describe the estimation framework in COXMEG using a penalized partial likelihood (PPL) proposed by (Breslow and Clayton, 1993; Ripatti and Palmgren, 2000) based on the Laplace approximation. Other estimation strategies such as the penalized likelihood method (McGilchrist, 1993; McGilchrist and Aisbett, 1991), the hierarchical likelihood (h-likelihood) framework (Ha et al., 2001, 2011; Lee et al., 2006), and the expectation–maximization (EM) algorithm (Cortiñas Abrahantes and Burzykowski, 2005) lead to a similar iterative procedure. Specifically, suppose that we have right-censored time-to-event observations *T*_*i*_,*i* ∈ {1,…,*N*} from a cohort of *N* subjects and a corresponding variable *d*_*i*_ indicating whether a failure (*d*_*i*_=1) or censoring (*d*_*i*_=0) occurs at *T*_*i*_ for each subject. The semiparametric proportional hazards (or Cox) model (Cox, 1972) assumes that the conditional hazard function for subject *i* has the form

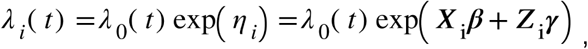

where *λ*_0_(*t*) is an unknown baseline hazard function, ***X*** is a matrix of *m* predictors corresponding to the fixed (***β*** ∈ ℜ^*m*×1^) effects, and ***Z*** is a design matrix for the random (***γ*** ∈ ℜ^*N*×1^) effects, respectively. The fixed effects include both the SNP to be tested and other covariates. In a CMEM, we assume that ***Z*** is the identity matrix (often denoted as ***I***), and ***γ*** follows a multivariate normal distribution, i.e.,

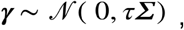

where *Σ* ∈ ℜ^*N*×*N*^ is expected to be a symmetric positive definite (SPD) matrix measuring the genetic distance between the subjects (e.g., a kinship matrix or a GRM), and *τ* is the variance of the genetic component. For simplicity, we consider here one variance component, but generalization to multiple components is straightforward. The situation where *Σ* is symmetric positive semidefinite (SPSD) is more complicated, and will be discussed in a separate section. The log-likelihood based on the above model is given by

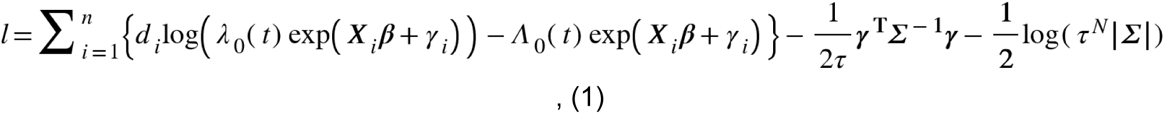

where 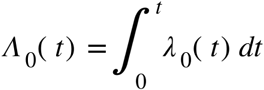 is the cumulative hazard function. The baseline hazard *λ*_0_(*t*) is unknown, but can be estimated from the data. Either substituting in e.q. (1) with a partial likelihood (McGilchrist, 1993; Ripatti and Palmgren, 2000) or using maximum likelihood estimation (MLE) for *λ*_0_(*t*) based on the h-likelihood (Ha et al., 2001) leads to the following PPL or equivalently h-likelihood

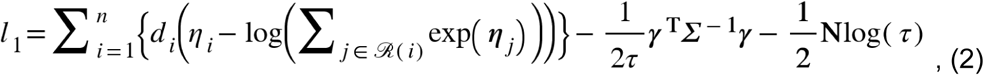

where *ℛ* (*i*) is the set of all subjects at risk when an event is observed for subject *i*.

Because ***β*** and ***γ*** are mutually canonical (see e.g., (Lee et al., 2006)), the h-likelihood *l*_1_ can be treated as an ordinary likelihood for estimating ***β*** and ***γ*** simultaneously, which we denote by ***θ*** = (***β, γ***), when *τ* is given (Ha et al., 2001; Lee et al., 2006). However, *τ* has to be estimated separately using a marginal likelihood or an adjusted profile likelihood. Following (Ripatti and Palmgren, 2000), an integrated marginal likelihood of *τ* using the Laplace approximation is

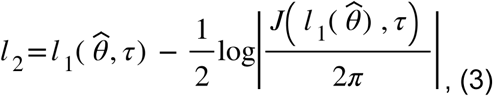

where *J* (*l*_*1*_,*τ*) is an information matrix of ***γ*** using the PPL *l*_*1*_ with 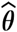 estimated from e.q. (2). A standard two-layer algorithm for estimating ***θ***, such as coxme, is to iterate between *l*_*1*_ and *l*_*2*_. In GWAS, however, it is often sensible to practically assume that *τ* is a constant for estimating the effect of each SNP (Kang et al., 2010; Loh et al., 2015). We leverage this assumption in COXMEG, which estimates the variance component *τ* under a null model with only relevant covariates, and then use the estimated 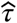 to analyze all SNPs. This step substantially reduces computational time compared to coxme, which estimates *τ* for each SNP individually. Some studies show that using a uniform variance component may lead to loss of statistical power to detect very significant SNPs (Zhou and Stephens, 2012). Nevertheless, this potential issue can be amended by conducting a sensitivity analysis for those SNPs passing certain significance in the first round of the GWAS. We first focus on algorithms for the maximization of *l*_*1*_, where an optimization algorithm finds 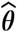 given a known 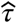. The estimation of *τ* under the null model will be discussed in a later section in detail.

### Estimation algorithm in COXMEG-sparse

We consider in COXMEG-sparse a situation where *Σ* is a sparse SPD matrix. Because in a sample of large size, *l*_*1*_ is highly quadratic in adjacent of the optimal ***θ***, the Newton-Raphson (NR) method generally converges in a small number of steps, and thus is an efficient optimization method for this high-dimensional problem. Computing the first and second derivatives *l*_*1*_ of with respect to ***θ*** gives the following score function and the Fisher information matrix

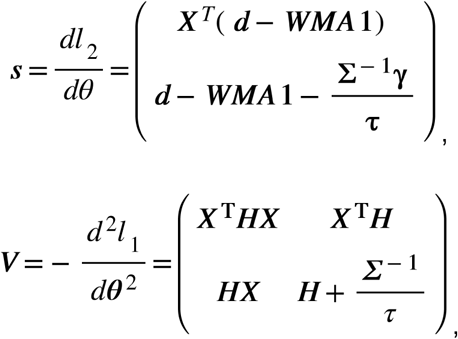

where

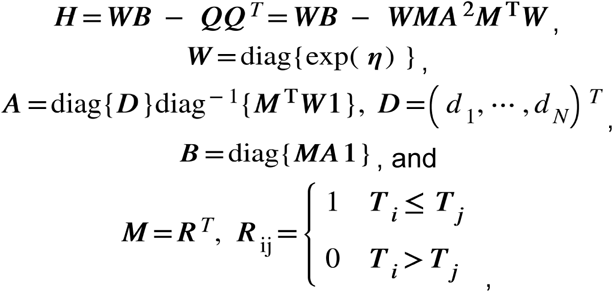

where *R* is an indicator matrix for the risk set *ℛ* (*i*), in which we adopt the Brelsow’s approximation (Breslow, 1974) for ties. The NR method estimates ***θ*** using

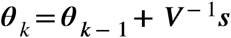

iteratively ***θ***_*k*_ until converges. In terms of computational burden, the cross product ***QQ***^*T*^ takes *O*(*N*^2^) rather than *O*(*N*^3^) because *W* and *A* are diagonal, and *M* is an indicator matrix similar, through permutation, to a triangular matrix when no ties are present. Another computational challenge is to compute ***V***^**−1**^***s***, which takes *O*(*N*^3^) if solved directly using e.g., Cholesky decomposition (Cholesky decomposition is valid because ***H*** is always SPSD as shown in Supplementary materials A.2, and ***V*** is SPD when is *Σ* SPD). If there are *p* SNPs and *L*_1_ iterations are needed on average in the NR optimization of *l*_1_ for each SNP, the total computational burden in the estimation of ***θ*** would be *O*(*N*^3^*pL*_1_), which becomes immediately intractable when the sample size *N* becomes thousands. We notice that when *Σ* is block-diagonal (e.g., a pedigree-based kinship matrix), coxme uses an approximated NR method by computing only the block-diagonal entries of ***V*** corresponding to those in *Σ*. Such an approximation reduces the computational time to at most *O*(*F*^2^*NpL*_1_), where *F* is the largest block (or family) size. In family-based studies, *F* is generally much smaller than *N*, and can often be assumed as a constant that does not increase with *N*. This approximation provides remarkable gain in computational performance. However, the drawback of this method is that this approximation depends on the structure of *Σ* and is limited to a block-diagonal *Σ*. Accordingly, more general GRMs with long-range correlations, e.g., long-distance cryptic relatedness, or other sparse correlation matrices are not efficiently handled in this framework. In addition, the block-diagonal approximation is not optimal when there are large sparse blocks in *Σ*. We next propose an algorithm which is efficient for any sparse SPD matrix.

Instead, we propose another efficient algorithm leveraging the sparsity. Rewriting ***V***^−1^using Schur complements leads to

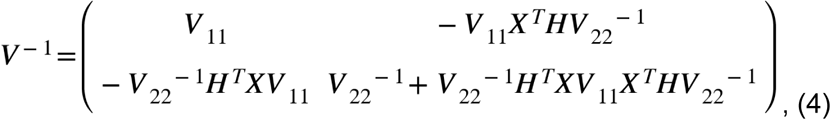

where

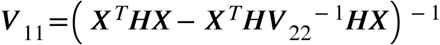

and

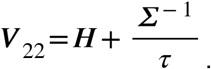

Note that the computation of ***X***^*T*^***H*** involves no explicit evaluation of ***QQ***^*T*^, and requires only multiplication and addition in *O*(*N*). Therefore, once ***V***_22_ is solved, the remaining computation can be done in *O*(*N*). The obstacle of inversing ***V***_22_ is that ***V***_22_ is fully dense even if *Σ* is sparse because ***QQ***^***T***^ is a fully dense matrix. To solve ***V***_22_, observing that ***WB*** is diagonal ***QQ***^*T*^, is low-rank, and *Σ* (or *Σ*^−1^) is sparse, rewriting ***V***_22_^−1^ using the Woodbury identity (Hager, 1989) gives

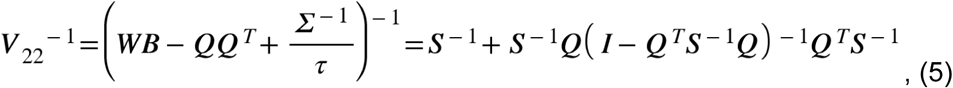

where

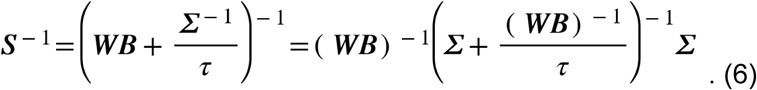

The advantage of using e.q. (5) is that it disentangles the sparse and dense parts in ***V***_22_^−1^, so that the dominant sparse term ***S***^−1^ in e.q. (6) can be evaluated efficiently by leveraging sparse algorithms depending on which of *Σ* and *Σ*^−1^ has better sparsity pattern. For example, when *Σ* is block-diagonal, both matrices are sparse. When *Σ* is a matrix with AR(1) correlation structure (Example 4 in (Liang and Zeger, 1986)), only *Σ*^−1^ is sparse. However, in many cases, *Σ*^−1^ might not be as sparse as *Σ*. The major computational bottleneck in e.q. (5) is the evaluation of (*I*−*Q*^*T*^*S*^−1^*Q*) ^−1^, which is at least *O*(*N*_1_^3^) where *N*_1_ is the number of non-censored subjects. So we simply ignore it in COXMEG-sparse, and propose to use instead the approximation

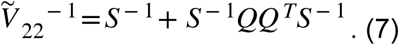

Substituting ***V***_22_^-1^ by 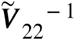 in e.q. (4) gives an approximated NR method. Similarly, no explicit evaluation of ***QQ***^*T*^ is needed when using e.q. (7). Hence, once a sparse Cholesky decomposition (SCD) of *S* is obtained, the evaluation of the second term in e.q. (7) mostly requires only one more forward-backward substitution. In the special case where *Σ* is block-diagonal, the computational burden for solving a linear system of 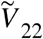 is *O*(*F*^2^*N*).

We show in Supplementary materials A.1 that 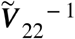 is a first-order approximation of ***V***_22_^-1^. In fact, using Neumann series, ***V***_22_^-1^ can be expressed as

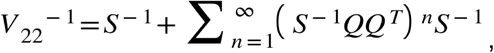

from which we see that *S*^-1^is a zero-order approximation of ***V***_22_^-1^, which is equivalent to completely ignoring -***QQ***^*T*^ from the information matrix ***V***. This amounts to adding ***QQ***^*T*^, a SPSD matrix, to the information matrix. We show in Supplementary materials A.2 that COXMEG-sparse satisfies the conditions for a class of inexact Newton methods (Dembo et al., 1982; Eisenstat and Walker, 1996). Intuitively, this suggests that this class of approximations still has the local convergence as long as a forcing term is bounded away from one (Dembo et al., 1982; Kelley, 1995) although it no longer has the quadratic convergence rate generally (i.e., it converges in more iterations than the NR method). We prove the convergence in more detail in Supplementary materials A.2, but in short, the convergence rate depends on multiple properties including the variance component *τ* and the spectrum density of *Σ*. In terms of computational efficiency, a higher-order approximation speeds up the convergence, which, on the other hand, requires one more forward-backward substitution. Therefore, there is a trade-off between the convergence rate and the number of forward-backward substitutions in selecting an optimal order of approximation. We propose the first-order approximation 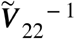 because our simulation and real studies showed that the first-order approximation is often adequate to achieve close to optimal performance in many practical applications to family-based GWAS (Figure S3, Supplementary materials A.2). In the cases where the algorithm converges slowly with 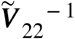, which often happens for a large *τ* or condition number of *Σ*, a higher-order approximation should be used for better computational performance (See additional results in Supplementary materials A.2). Finally, replacing ***V***_22_^-1^ by 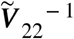 in *V*_11_ gives the following approximated variance of 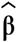

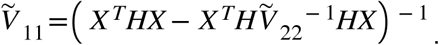

We found that the relative error between 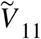 and *V*_11_ was <0.1% under most settings considered in our simulation except for *τ*=0.05 (Figure S7), suggesting that the first-order approximation 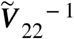 is sufficiently accurate for testing ***β*** in general applications in GWAS when the variance component *τ* is not overly large. Otherwise, the second-order approximation should be used.

As aforementioned, the major step in evaluating 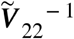 is to compute *S*^−1^. Either term in e.q. (6) can be used depending on which matrix, *Σ* or *Σ*^−1^, has more sparsity. In addition, certain SCD algorithms may produce fewer fill-ins, and thus one or another algorithm can be faster. In our simulations, we compared three sparse algorithms in R to solve the linear system of *S*, (i) the LDLT SCD in RcppEigen, (ii) the sparse preconditioned conjugate gradient (PCG) using diagonal elements in RcppEigen, and (iii) the SCD in the Matrix R package. We found in our study that the LDLT SCD in RcppEigen yields the best performance under the vast majority of scenarios, especially for very sparse *Σ* (Figure S4). On the other hand, the PCG method shows computational advantages when *Σ* is very large or less sparse (Figure S4).

### Score test for dense relatedness matrices

When both the matrices *Σ* and *Σ* ^−1^ are dense, COXMEG-sparse can no longer leverage the sparse algorithms to speed up the evaluation of ***S***^−1^. Instead of the Cholesky decomposition used in coxme, we implemented a PCG method (Lidauer et al., 1999; Tsuruta et al., 2001) (termed as COXMEG-pcg) to compute each step in the NR algorithm. This alteration reduces the time complexity from to *O*(*N*^3^*pL*_1_) to *O*(*N*^2^*pL*_1_). However, for a large-scale GWAS, the PCG method might still be overly time-consuming. Therefore, we further propose COXMEG-score, a fast score test for the situation in which *Σ* and *Σ* ^−1^ cannot be approximated by a sparse matrix. In COXMEG-score, we first fit the full CMEM under the null model, in which only covariates are included in *X*. After the null model is fitted, we obtain the estimates of the score function of ***γ***, ***H***, and the inverse of the information matrix under the null hypothesis, denoted by ***s***_0_, ***H*** _0_ and ***V*** _0_^−1^, respectively, where ***s***_0_ =***d*** − ***Q*** · 1. Denote by ***z*** a vector of the genotypes of a SNP to be tested. The score test statistic proposed in COXMEG-score is given by

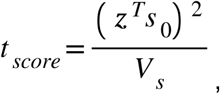

where ***V***_*s*_ is the variance of the score in the numerator given by

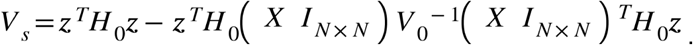

Because the h-likelihood *l*_1_ can be treated as an ordinary likelihood once the variance component is given, *t*_*score*_ approximately follows a χ_1_^2^ distribution with one degree of freedom under the null hypothesis that the SNP has no effect on the HR. Note that the computation of *V*_*s*_ involves only one matrix-vector multiplication and the rest can be performed in *O*(*N*). So the computational burden of the score test alone is *O*(*N*^2^_*P*_) regardless of the structure of the relatedness matrix. This time complexity is the same as the score tests proposed for generalized linear models (Chen et al., 2016, 2019). Our simulation results suggest that COXMEG-score is statistically as powerful as the Wald tests like COXMEG-sparse and coxme (Figures S1, S2).

### Estimation of variance components

We have described above the estimation and testing algorithms when the variance component τ is given. Here, we describe the estimation of *τ* In COXMEG, *τ* is estimated only once under the null model, and then is used for testing all SNPs. The estimation of can be performed by calculating the first and second derivatives of *l*_2_ and iterating between *l*_1_ and *l*_2_ using the NR method as suggested in (Ha et al., 2011; Ripatti and Palmgren, 2000). We found in practice that this EM-like procedure converges rather slowly for estimating *τ*. Instead, we follow the strategy implemented in coxme, which treats *l*_2_ as a marginal likelihood of *τ* and directly uses a derivative-free optimization because is often a scalar or in low-dimension. We found in our simulation study that this strategy converges faster. Thus, after *l*_1_ is optimized, the only additional computation in each iteration for optimizing *l*_2_ is the evaluation of the log-determinant of *J*(*l*_1_, *τ*)=*V*_22_. As *V*_22_ is SPD, a direct evaluation would require a Cholesky decomposition, whose time complexity is 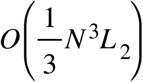, where *L*_2_ is the number of iterations in optimizing *l*_2_ which is often between 10 and 50 depending on the convergence. We evaluated the performance of the following three derivative-free algorithms, (i) Bobyqa (Powell, 2009) implemented in the nloptr R package (Ypma, 2014), (iii) Brent’s method and (iii) the Nelder-Mead method (Nelder and Mead, 1965) in the optim R function. We found in our simulation studies that Bobyqa is often the fastest, and Brent’s method is slightly slower than Bobyqa.

When *N* is very large (e.g., >20,000), the evaluation of the log-determinant becomes a bottleneck in COXMEG-sparse and COXMEG-score, and even intractable. In addition, a second-stage sensitivity analysis using a variant-specific variance component may be performed for those SNPs passing a predefined threshold of significance in the first-round. When the number of significant SNPs is large, this second-stage analysis would be very computational intensive. So, we propose fast approximate methods to estimate *τ* for sparse and dense GRMs, respectively. Specifically, for a sparse GRM, we propose the following marginal likelihood instead of *l*_2_

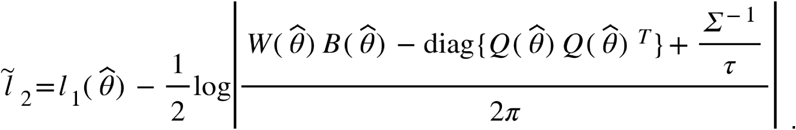

Compared to *l*_2,_ the only difference is that 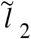 uses a diagonal approximation of ***QQ***^*T*^. When *Σ* or *Σ* ^−1^ is sparse, the determinant can often be efficiently computed using a SCD depending on its sparsity pattern. We found in our simulation study that the estimated *τ* using 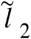 is very close to that from coxme, which uses a block-diagonal approximation of ***QQ***^*T*^, with a relative difference <1% in most cases (Figure S5).

When the relatedness matrix is dense or its sparsity pattern results in too many fill-ins, the above diagonal approximation does not help. Instead, we propose a SLQ method (Bai et al., 1996; Golub and Meurant, 2009; Ubaru et al., 2017) to approximate the log-determinant, which reduces the time complexity from cubic to quadratic (*O*(*n*_*m*_*n*_*q*_*N*^2^*L*_2_), see Supplementary materials A.4). The basic idea is to first express the log-determinant using a Monte Carlo estimator of the trace of the matrix logarithm. Each term in the Monte Carlo estimator can be written as a Riemann-Stieltjes integral approximated by the Gaussian quadrature rule. The nodes and weights in the Gaussian quadrature can be elegantly computed by the eigenvalues and first elements in the eigenvectors of the tridiagonal matrix obtained from the Lanczos algorithm. We found in our simulation study that the accuracy of the SLQ method for approximating a log-determinant was generally higher by one order of magnitude than that using Chebyshev orthogonal polynomials (Han et al., 2016; Pace and LeSage, 2004), both of which were more accurate than Martin’s Taylor expansion (Barry and Kelley Pace, 1999; Martin, 1992), consistent with previous reports (Han et al., 2016; Ubaru et al., 2017). More details about the SLQ approximation were described in Supplementary materials A.4.

### Handling a positive semidefinite relatedness matrix

When the relatedness matrix *Σ* is SPSD, *l*_1_ becomes invalid because *Σ*^−1^ does not exist. So, we instead propose a generalized penalized partial likelihood (GPPL)

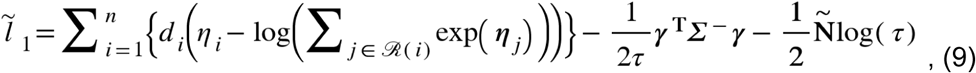

where *Σ* ^−^ is the pseudoinverse defined as *Σ* ^−^,=*U Λ* ^−^*U*^*T*^ where *U* is the eigenvectors of *Σ*, and *Λ*^−^ is the pseudoinverse of the diagonal matrix consisting of the eigenvalues of *Σ*. In addition, *N* is replaced by 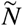, the number of non-zero eigenvalues of *Σ*. The rationale of e.q. (9) is similar to that in (He et al., 2016c; Kauermann and Wegener, 2011), in which the penalty is imposed on a subspace of the coefficients. We show in Supplementary materials A.3 that when replacing *l*_1_ with the GPPL 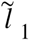 for a SPSD *Σ*, the NR algorithms in COXMEG-sparse and COXMEG-score still work except that the sum of elements in each row of *Σ* is zero (i.e., an SPSD matrix has **1** eigenvector with eigenvalue 0). In such case,*V*_22_ becomes non-invertible. In addition, the last transformation in e.q. (6) is valid only for *Σ* being SPD. So, if *Σ* but not *Σ* ^−1^ is sparse, it would be better to make *Σ* SPD, as discussed in the following section.

### Construction of relatedness matices for COXMEG

Depending on the purpose of using the CMEM, the relatedness matrix *Σ* summarise the dependence structure between individuals. Such a matrix in the context of GWAS is generally a kinship matrix or a GRM. There exist a variety of approaches defining and calculating these matrices depending on the assumption of the genetic architecture underlying the phenotype. Here, we focus on two frequently used types of relatedness matrices, a pedigree-based kinship matrix and a GCTA GRM based on the infinitesimal model (Yang et al., 2011) and discuss how to construct such matrices for COXMEG.

A pedigree-based kinship matrix provides a measure of the total average genetic correlation based on the pedigree information but no genotype information, and it does not account for a specific phenotype. When the individuals in a cohort are more or less ethnically homogeneous, the CMEM with a pedigree-based kinship matrix is used primarily to account for family structure. A pedigree-based kinship matrix (for example, generated by the kinship2 R package (Sinnwell et al., 2014)), is generally block-diagonal SPD in which individuals from different families are regarded as independent. One exception is MZ twins, which lead to the kinship matrix being SPSD. COXMEG-sparse is suitable for this situation because such matrices are sparse when the family size is not at the same magnitude of the sample size.

In contrast, a GRM is more flexible in terms of genomic regions, more precise in terms of measuring genetic correlation, and can detect long-distance cryptic relatedness. A GCTA GRM based on the infinitesimal model (Yang et al., 2011) is calculated by

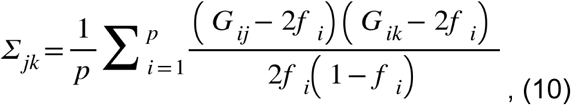

where *f* _*i*_ is the frequency of the coding allele of SNP *i* and *G*_*ij*_ is the dosage of SNP *i* and individual *j*. This type of GRMs is a covariance matrix, which is generally SPSD even if *p* > *N* because one degree of freedom is lost for estimating *f* _*i*_ from the same sample. It always has eigenvector **1** with eigenvalue 0. Note that an estimated GRM may have negative eigenvalues if the SNPs used to construct it have many missing values, which should be avoided.

When the sample has both family structure and population stratification, the GCTA GRM is not sparse. The raw GCTA GRM cannot be directly used for COXMEG because it is SPSD and always has eigenvector **1** with eigenvalue 0. Nevertheless, this is not a big problem in most cases because the actual sample size included in the analysis is often smaller than the total sample size in the relatedness matrix due to removal of individuals with missing data or early censored. Otherwise, a scaled GRM, in which the diagonal elements are all one, is always valid for COXMEG. In addition, for an ethnically homogeneous cohort, a GRM can often be approximated by a sparse matrix using thresholding because most elements outside of a family are almost zero. Various methods can be used to approximate the GRM, such as hard thresholding (Bickel and Levina, 2008), and soft thresholding using graphical lasso (Friedman et al., 2008). We found in our analysis of the NIA-LOADFS data that simple hard thresholding, which has literally little computational cost, using a small cutoff (e.g., 0.02) could induce sparsity and often (but not always) preserve SPD (Bickel and Levina, 2008). Sparsity approximation with guaranteed positive definite is also available (e.g., (Rothman, 2012) implemented in the PDSCE R package) albeit much computational intensive compared to a simple hard thresholding.

### Simulation study

We performed simulation studies to investigate the computational and estimation performance under R 3.5. The simulations for computational performance were conducted on Windows using a PC with Intel Core i7-8700. Some simulations for evaluating the estimation performance were conducted on Linux. We considered block-diagonal correlation matrices, banded AR(1) correlation matrices, sparse GRMs by thresholding a GRM estimated from the genotype data in the NIA-LOADFS. All blocks in the block-diagonal correlation matrices share the same size and a single correlation coefficient. We evaluated block sizes of 5, 20, 100, 500, and correlation coefficients of 0.1, 0.5, and 0.9. The elements in the banded AR(1) correlation matrices were generated by *ρ* ^|*i* −*j* |^ and then set zeros for all *ρ* ^|*i* −*j* |^ < 0.01, in which we evaluated *ρ*=0.3,0.6,0.9. For each relatedness matrix *Σ*, we generated random effects based on ***γ*** ∼ *Σ* (0, *τ Σ*), in which we considered *τ*=0.01,0.05,0.1,0.2,0.5 to reflect different magnitudes of heritability. Genotype dosages for each SNP were generated from a binomial distribution *G∼B*(2,*p*), in which *p*, the MAF, ranged between 0.05 and 0.5. A disease time variable was generated using the following exponential distribution ***Y****∼Exp* (exp(*μ*+***G***+ ***γ***)), in which we assumed a constant baseline hazard with *μ*= −2. For each individual, we generated a censoring time variable using the following exponential distribution *C*∼*Exp*(*c*), in which we considered *c*=0.02,0.05,0.1,0.15 to reflect scenarios from heavy censoring *c*=0.15 to almost no censoring *c* = 0.02. The time-to-event variable was generated by

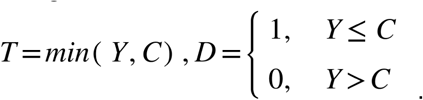

## Supporting information

R_package_vignettes

## Acknowledgements

This manuscript was prepared using limited access datasets obtained though dbGaP (accession numbers: phs000168.v2.p2 (LOADFS). This research was supported by Grants from the National Institute on Aging (P01 AG043352, R01 AG047310 and R01 AG061853). The funders had no role in study design, data collection and analysis, decision to publish, or manuscript preparation. The content is solely the responsibility of the authors and does not necessarily represent the official views of the National Institutes of Health.

## Author contributions

LH conceived, developed and implemented the algorithms. LH analyzed the simulated and real data. AK contributed to acquiring the data, examining the algorithms and discussion of final results. LH and AK contributed to the writing of the manuscript.

## Supplementary Tables

Table S1. Results of 23 common SNPs identified by a recent meta-analysis of AD. The p-values of HRs were obtained from COXMEG-score based on a score test. The position of the SNPs is based on hg19.

## Supplementary Figures

Figure S1. Comparison of empirical FPR of coxme, COXMEG-sparse, COXMEG-score, and coxph with a shared frailty. The relatedness matrix used in the simulation is a block-diagonal correlation matrix with the block size ranging between 5-100. We evaluated the FPR for the correlation ρ in each block being 0.1, 0.5, and 0.9, and the variance component *τ* between 0.02 and 0.5.

Figure S2. Comparison of empirical power of coxme, COXMEG-sparse, COXMEG-score, and coxph with a shared frailty. The relatedness matrix used in the simulation is a block-diagonal correlation matrix with the block size ranging between 5-100 and the correlation ρ ranging between 0.1 and 0.9. We evaluated the power for the HRs being 0.01, 0.05, and 0.1, and the sample size between 1000 and 5000. We also assess the power under no censoring, moderate censoring and heavy censoring.

Figure S3. Evaluation of the convergence rate of higher-order approximations in COXMEG-sparse. The relatedness matrix used in the simulation is a block-diagonal correlation matrix with the block size ranging between 5-100 and the correlation ρ ranging between 0.1 and 0.9. For each setting, the convergence rate was measured by the time for estimating the HRs of one predictor given a variance component *τ* ranging from 0.02 to 0.5.

Figure S4. Evaluation of the computational performance of three methods (RcppEigen::LDLT, Matrix::solve using the Cholesky decomposition, and RcppEigen::CG with diagonal preconditioned) for solving the sparse linear system in COXMEG-sparse. The relatedness matrix used in the simulation is a block-diagonal correlation matrix with the block size varying between 5-500 and the correlation ρ being 0.5. For each setting, the convergence rate was measured by the time for estimating the HRs of one predictor given a variance component *τ* ranging from 0.02 to 0.5.

Figure S5. Comparison of estimated variance components by coxme, COXMEG-sparse with the exact log-determinant, and COXMEG-sparse with a diagonal approximated log-determinant. (A) COXMEG-sparse with a diagonal approximated log-determinant vs. coxme; (B) COXMEG-sparse with the exact log-determinant vs. coxme; (C) COXMEG-sparse with a diagonal approximated log-determinant vs. the exact log-determinant.

Figure S6. Comparison of estimated variance components by COXMEG-score with the exact log-determinant, and COXMEG-score with the SLQ approximation under sample sizes of 5000 and 10,000. The relatedness matrix used in the simulation is a block-diagonal correlation matrix with the block size varying between 5-500 and the correlation ρ between 0.1 and 0.9.

Figure S7. Relative error between the variance of log(HR) estimated using an approximated *V*_22_^−1^ in COXMEG-sparse and using an exact Hessian matrix. Four approximations of *V*_22_^−1^ (Zero-order to Third-order) were evaluated under settings of different sample sizes, and variance components. The red dots are the mean values.

