## Supplementary materials for "Genome-wide association analysis of age-at-onset traits using Cox mixed-effects models"

### Supplementary materials for "Genome-wide association analysis of age-at-onset traits using Cox mixed effects model"

**A. Supplementary text** **1**
1. Proof that is the first-order approximation of in COXMEG-sparse 1
2. Local convergence of COXMEG-sparse 3
3. Proof of the validity of the NR method for positive semidefinite covariance matrices 6
4. Approximation of the log-determinant using the SLQ method 7
5. Simulation study for statistical power and empirical size 9

**B. Supplementary Tables** **9**

**C. Supplementary Figures** **11**

**References** **18**

### A. Supplementary text

#### 1. Proof that $\tilde{V}_{22}^{-1}$ is the first-order approximation of $V_{22}^{-1}$ in COXMEG-sparse

Here, we show that $\tilde{V}_{22}^{-1} = S^{-1} + S^{-1}QQ^TS^{-1}$ is a first-order approximation of $\boldsymbol{V}_{22}^{-1}$, where $S^{-1} = \left(WB + \frac{\Sigma^{-1}}{\tau}\right)^{-1}$. We follow the notations used in the main text. One useful observation about $QQ^T$ is that the columns corresponding to the censored subjects in $\boldsymbol{M}$ do not contribute to $QQ^T$ because their corresponding elements in $\boldsymbol{A}$ are zero. Denote by $\tilde{\boldsymbol{M}} \in \{0, 1\}^{N \times N_1}$ the matrix after removing the columns corresponding to the censored subjects from $\boldsymbol{M}$, where $N_1$ is the number of subjects experiencing the event of interest. We rewrite $H$ as

$$
\begin{aligned}
\boldsymbol{H} &= \boldsymbol{W}\boldsymbol{B} - \boldsymbol{Q}\boldsymbol{Q}^T \\
&= \boldsymbol{W}\boldsymbol{B} - \boldsymbol{W}\boldsymbol{M}\boldsymbol{A}^2\boldsymbol{M}^{\mathrm{T}}\boldsymbol{W} \\
&= \boldsymbol{W}\boldsymbol{B} - \boldsymbol{W}\tilde{\boldsymbol{M}}\tilde{\boldsymbol{A}}^2\tilde{\boldsymbol{M}}^T\boldsymbol{W} \\
&= \boldsymbol{W}\boldsymbol{B} - \tilde{\boldsymbol{Q}}\tilde{\boldsymbol{Q}}^{\mathrm{T}}
\end{aligned}
\quad , \text{(S1)}
$$

where

$$
\tilde{\boldsymbol{Q}} = \boldsymbol{W}\tilde{\boldsymbol{M}}\tilde{\boldsymbol{A}}
$$

$$
\tilde{\boldsymbol{A}} = \mathrm{diag}^{-1}\{\tilde{\boldsymbol{M}}^T\boldsymbol{W}\mathbf{1}\}, \text{(S2)}
$$

$$
\boldsymbol{B} = \mathrm{diag}\{\boldsymbol{M}\boldsymbol{A}\mathbf{1}\} = \mathrm{diag}\{\tilde{\boldsymbol{M}}\tilde{\boldsymbol{A}}\mathbf{1}\}. \text{ (S3)}
$$

Because $\boldsymbol{S}^{-1}$ is always positive definite after removing those subjects censored before the first failure, we then rewrite ${\boldsymbol{V}_{22}}^{-1}$ in the following way

$$
\begin{aligned}
&{\boldsymbol{V}_{22}}^{-1} \\
&= \left(\boldsymbol{W}\boldsymbol{B} - \boldsymbol{Q}\boldsymbol{Q}^T + \frac{\boldsymbol{\Sigma}^{-1}}{\tau}\right)^{-1} \\
&= \left(\left(\boldsymbol{W}\boldsymbol{B} + \frac{\boldsymbol{\Sigma}^{-1}}{\tau}\right)\left(\boldsymbol{I} - \left(\boldsymbol{W}\boldsymbol{B} + \frac{\boldsymbol{\Sigma}^{-1}}{\tau}\right)^{-1}\boldsymbol{Q}\boldsymbol{Q}^T\right)\right)^{-1} \\
&= \left(\boldsymbol{I} - \boldsymbol{S}^{-1}\boldsymbol{Q}\boldsymbol{Q}^T\right)^{-1}\boldsymbol{S}^{-1}
\end{aligned}
\quad , \text{ (S4)}
$$

Denote by $\|\boldsymbol{A}\|$ the spectral norm of $\boldsymbol{A}$, which is the largest eigenvalue of $\boldsymbol{A}$ when $\boldsymbol{A}$ is SPD (i.e., $\|\boldsymbol{A}\|$ is $\lambda_{max}(A)$ ). Now we show that the spectral norm of $\boldsymbol{S}^{-1}\boldsymbol{Q}\boldsymbol{Q}^T$ in the last line in e.q. (S4) is strictly less than one, that is,

$$
\left\|\boldsymbol{S}^{-1}\boldsymbol{Q}\boldsymbol{Q}^T\right\| < 1 .
$$

Note that both $V_{22}$ and $\boldsymbol{S}^{-1}$ are SPD, so their product

$$
\begin{aligned}
&\boldsymbol{S}^{-1}\boldsymbol{V}_{22} \\
&= \boldsymbol{S}^{-1}(\boldsymbol{S} - \boldsymbol{Q}\boldsymbol{Q}^T) \\
&= \boldsymbol{I} - \boldsymbol{S}^{-1}\boldsymbol{Q}\boldsymbol{Q}^T
\end{aligned}
$$

has only positive eigenvalues. This implies that all eigenvalues of $\boldsymbol{S}^{-1}\boldsymbol{Q}\boldsymbol{Q}^T$ are strictly less than one. Therefore, we can rewrite ${\boldsymbol{V}_{22}}^{-1}$ by expanding the first inverse term in e.q. (S4) using a Neumann series, which gives

$$
\begin{aligned}
&{\boldsymbol{V}_{22}}^{-1} \\
&= \left(\boldsymbol{I} - \boldsymbol{S}^{-1}\boldsymbol{Q}\boldsymbol{Q}^T\right)^{-1}\boldsymbol{S}^{-1} \\
&= \left(\boldsymbol{I} + \boldsymbol{S}^{-1}\boldsymbol{Q}\boldsymbol{Q}^T + \boldsymbol{S}^{-1}\boldsymbol{Q}\boldsymbol{Q}^T\boldsymbol{S}^{-1}\boldsymbol{Q}\boldsymbol{Q}^T + \cdots\right)\boldsymbol{S}^{-1} \\
&= \sum_{k=0}^{\infty}\boldsymbol{S}^{-1}\left(\boldsymbol{Q}\boldsymbol{Q}^T\boldsymbol{S}^{-1}\right)^k
\end{aligned}
\quad ,
$$

from which we obtain ${\tilde{\boldsymbol{V}}_{22}}^{-1}$ by taking the first two terms in the series.

#### 2. Local convergence of COXMEG-sparse

We first show that COXMEG-sparse is locally convergent, and then discuss the factors affecting its convergence rate. There are multiple ways to investigate the local convergence of COXMEG-sparse. Our approach is to show that COXMEG-sparse belongs to a class of inexact Newton methods (Dembo et al., 1982) with the following property

$$\left\| -V(\boldsymbol{\theta}_c)\,\boldsymbol{e} + \boldsymbol{s}(\boldsymbol{\theta}_c) \right\| \le \eta_c \left\| \boldsymbol{s}(\boldsymbol{\theta}_c) \right\|, \quad \text{(S5)}$$

where $\boldsymbol{e}$ is the step change in each iteration in COXMEG-sparse, and $\boldsymbol{s}(\boldsymbol{\theta}_c)$ and $V(\boldsymbol{\theta}_c)$ are the score function and negative Hessian evaluated at the current step $\boldsymbol{\theta}_c$, respectively. The goal is to show that the step $\boldsymbol{e}$ chosen in COXMEG-sparse satisfies inequality (S5) with the forcing term $\eta_c < 1$. We first consider the convergence of using the zero-order approximation of $\boldsymbol{V}_{22}{}^{-1}$ (i.e., replacing $\boldsymbol{V}_{22}{}^{-1}$ by $\boldsymbol{S}^{-1}$ in $\boldsymbol{V}^{-1}$ ). Note that the zero-order approximation of $\boldsymbol{V}_{22}{}^{-1}$ amounts to adding $\boldsymbol{QQ}^T$ to the bottom-right corner of $\boldsymbol{V}$. So, the iteration step in this case is

$$\boldsymbol{e} = \tilde{\boldsymbol{V}}^{-1}(\theta_c)\, s(\theta_c) = \left( V(\theta_c) + \boldsymbol{Q}_0 \right)^{-1} s(\theta_c), \quad \text{(S6)}$$

where we denote $\boldsymbol{Q}_0 = \begin{pmatrix} 0 & 0 \\ 0 & \boldsymbol{QQ}^T \end{pmatrix}$. Plugging e.q. (S6) into the left-hand side of (S5) gives

$$\begin{aligned}
&\left\| -V(\theta_c)\,\boldsymbol{e} + s(\theta_c) \right\| = \left\| -V(\theta_c)\left( V(\theta_c) + \boldsymbol{Q}_0 \right)^{-1} s(\theta_c) + s(\theta_c) \right\| \\
&= \left\| \left( \boldsymbol{I} - V(\theta_c)\left( V(\theta_c) + \boldsymbol{Q}_0 \right)^{-1} \right) s(\theta_c) \right\| \\
&\le \left\| \left( \boldsymbol{I} - V(\theta_c)\left( V(\theta_c) + \boldsymbol{Q}_0 \right)^{-1} \right) \right\| \left\| s(\theta_c) \right\| \\
&= \left\| \boldsymbol{Q}_0 \left( V(\theta_c) + \boldsymbol{Q}_0 \right)^{-1} \right\| \left\| s(\theta_c) \right\| \\
&= \eta_c \left\| s(\theta_c) \right\|
\end{aligned}$$

.

It remains to show that the forcing term $\eta_c$ is uniformly less than one. Under regularity conditions (e.g., Assumption 2.2.1 in (Kelley, 1999)), $\boldsymbol{V}(\theta_c)$ is Lipschitz continuous within a neighbourhood of $\theta_*$ and is nonsingular at $\theta_*$ for which $s(\boldsymbol{\theta}_*) = \boldsymbol{0}$. This suggests that we can always find a neighbourhood of $\theta_*$ so that the smallest eigenvalue of $\boldsymbol{V}(\theta_c)$ is bounded away from zero. Because the largest eigenvalue of $\boldsymbol{Q}_0$ is one, we can find a neighbourhood of $\theta_*$, within which $\boldsymbol{V}(\theta_c)\left( \boldsymbol{V}(\boldsymbol{\theta}_c) + \boldsymbol{Q}_0 \right)^{-1}$ has all positive eigenvalues bounded away from zero, that is,

$$\lambda_{min}\left( \boldsymbol{V}(\boldsymbol{\theta}_c)\left( \boldsymbol{V}(\boldsymbol{\theta}_c) + \boldsymbol{Q}_0 \right)^{-1} \right) \ge \varepsilon > 0, \quad \boldsymbol{\theta}_c \in \delta(\boldsymbol{\theta}_*)$$

where $\lambda_{min}(\cdot)$ denotes the smallest eigenvalue of a matrix. This implies

$$
\begin{aligned}
&\lambda_{min}\left( \boldsymbol{V}\left( \boldsymbol{\theta}_c\right) \left( \boldsymbol{V}\left( \boldsymbol{\theta}_c\right) + \boldsymbol{Q}_0\right)^{-1}\right) \\
&=\lambda_{min}\left( \left( \boldsymbol{V}\left( \boldsymbol{\theta}_c\right) + \boldsymbol{Q}_0 - \boldsymbol{Q}_0\right) \left( \boldsymbol{V}\left( \boldsymbol{\theta}_c\right) + \boldsymbol{Q}_0\right)^{-1}\right) \\
&=\lambda_{min}\left( \boldsymbol{I} - \boldsymbol{Q}_0\left( \boldsymbol{V}\left( \boldsymbol{\theta}_c\right) + \boldsymbol{Q}_0\right)^{-1}\right) \geq \varepsilon > 0 \\
&\Leftrightarrow 1 - \lambda_{max}\left( \boldsymbol{Q}_0\left( \boldsymbol{V}\left( \boldsymbol{\theta}_c\right) + \boldsymbol{Q}_0\right)^{-1}\right) \geq \varepsilon \\
&\Leftrightarrow \left\| \boldsymbol{Q}_0\left( \boldsymbol{V}\left( \boldsymbol{\theta}_c\right) + \boldsymbol{Q}_0\right)^{-1}\right\| \leq 1 - \varepsilon < 1
\end{aligned}
.
$$

Therefore, according to Theorem 2.3 in (Dembo et al., 1982), the algorithm is locally convergent at least linearly in the norm $\|\theta\|_* = \|V^{-1}\theta\|$ with asymptotic rate constant no greater than $1-\varepsilon$. When a higher-order approximation is used in COXMEG-sparse, the difference is to replace $QQ^T$ in e.q. (S6) by another positive semidefinite matrix with a smaller matrix norm. Therefore, the proof of the local convergence still follows.

The convergence rate is controlled by $\varepsilon$. If $\varepsilon$ is equal to 1, in which case no approximation of $\boldsymbol{V}_{22}^{-1}$ is used, the convergence rate becomes quadratic. Therefore, the convergence rate depends on how close the approximation $\tilde{\boldsymbol{V}}_{22}^{-1}$ is to $\boldsymbol{V}_{22}^{-1}$. We then assess which factors affect the convergence rate practically. Note that $\boldsymbol{V}_{22}^{-1}$ in e.q. (S4) can be written as

$$
\begin{aligned}
&\boldsymbol{V}_{22}^{-1} \\
&=\left( \boldsymbol{I} - \boldsymbol{S}^{-1}\boldsymbol{Q}\boldsymbol{Q}^T\right)^{-1}\boldsymbol{S}^{-1} \\
&=\left( \boldsymbol{I} - \left( \boldsymbol{W}\boldsymbol{B}\left( \boldsymbol{I} + (\boldsymbol{W}\boldsymbol{B})^{-1}\frac{\boldsymbol{\Sigma}^{-1}}{\tau}\right)\right)^{-1}\boldsymbol{Q}\boldsymbol{Q}^T\right)^{-1}\boldsymbol{S}^{-1} \\
&=\left( \boldsymbol{I} - \left( \boldsymbol{I} + (\boldsymbol{W}\boldsymbol{B})^{-1}\frac{\boldsymbol{\Sigma}^{-1}}{\tau}\right)^{-1}\boldsymbol{B}^{-1}\boldsymbol{W}^{-1}\boldsymbol{Q}\boldsymbol{Q}^T\right)^{-1}\boldsymbol{S}^{-1}
\end{aligned}
, \text{(S7)}
$$

where the inverses are valid because of SPD of $W$, $B$ and $\boldsymbol{WB}$ after removing all censored samples before the first occurrence of the event of interest. In COXMEG-sparse, we use a Neumann series to approximate the first inverse in (S7), so a lower-order approximation would have poor performance, at least in some direction, if the spectral norm of

$$
\left( \boldsymbol{I} + (\boldsymbol{W}\boldsymbol{B})^{-1}\frac{\boldsymbol{\Sigma}^{-1}}{\tau}\right)^{-1}\boldsymbol{B}^{-1}\boldsymbol{W}^{-1}\boldsymbol{Q}\boldsymbol{Q}^T \quad \text{(S8)}
$$

is close to one. To investigate the spectral norm of (S8), we substitute e.q. (S1), (S2) and (S3) into $QQ^T$ in (S8), which gives

$$
\begin{aligned}
&\left(I+(WB)^{-1}\frac{\Sigma^{-1}}{\tau}\right)^{-1}B^{-1}W^{-1}QQ^{T}\\
&=\left(I+(WB)^{-1}\frac{\Sigma^{-1}}{\tau}\right)^{-1}B^{-1}\tilde{\mathbf{M}}\tilde{\mathbf{A}}\tilde{\mathbf{Q}}^{T}\\
&=\left(I+(WB)^{-1}\frac{\Sigma^{-1}}{\tau}\right)^{-1}\underbrace{\mathrm{diag}^{-1}\{\tilde{\mathbf{M}}\tilde{\mathbf{A}}1\}\tilde{\mathbf{M}}\tilde{\mathbf{A}}}_{S_I}\underbrace{\mathrm{diag}^{-1}\{\tilde{\mathbf{M}}^{\mathrm{T}}W1\}\tilde{\mathbf{M}}^{T}W}_{S_{II}}
\end{aligned}
$$

It is clear that $S_I$ and $S_{II}$ are recognized as two rectangular row stochastic (also called Markov) matrices. We show that $\|S_I S_{II}\|=1$ (i.e., the spectral norm of $S_I S_{II}$ is one), and all of its eigenvalues are between zero and one. First, we can easily verify that $1$ is an eigenvector of $S_I S_{II}$ with eigenvalue one. Then, suppose that $\nu$ is an eigenvector of $S_I S_{II}$, and $\nu_{\mathrm{k}}$ is the element that has the largest absolute value of the non-zero elements in $\nu$, we have

$$
\begin{aligned}
&|\lambda\nu_{\mathrm{k}}|=\left|(S_I S_{II}\nu)_k\right|\\
&=\left|\nu_1\sum_i s_{Iki}s_{IIi1}+\nu_2\sum_i s_{Iki}s_{IIi2}+\ldots+\nu_N\sum_i s_{Iki}s_{IIiN}\right|\\
&\le\left|\sum_i s_{Iki}s_{IIi1}+\sum_i s_{Iki}s_{IIi2}+\ldots+\sum_i s_{Iki}s_{IIiN}\right||\nu_k|\\
&=\left|\sum_j\sum_i s_{Iki}s_{IIij}\right||\nu_k|\\
&=\left|\sum_i s_{Iki}\sum_j s_{IIij}\right||\nu_k|\\
&=\left|\sum_i s_{Iki}\right||\nu_k|\\
&=|\nu_k|
\end{aligned},
$$

from which we obtain $|\lambda|\le 1$ (i.e., all eigenvalues have its absolute value no larger than one).

Because $WB$ is positive definite, $S_I S_{II}$ is similar to $(WB)^{-\frac{1}{2}}QQ^T(WB)^{-\frac{1}{2}}$, which means that all eigenvalues of $S_I S_{II}$ are non-negative. Summarizing all evidence above implies that the eigenvalues of $S_I S_{II}$ are between zero and one. Next, we show that all eigenvalues of $\left(I+(WB)^{-1}\frac{\Sigma^{-1}}{\tau}\right)^{-1}$ are less than one. In fact, because $WB$ and $\Sigma^{-1}$ are SPD, $(WB)^{-1}\Sigma^{-1}$ is similar to $(WB)^{-\frac{1}{2}}\Sigma^{-1}(WB)^{-\frac{1}{2}}$, which is a quadratic form and has all positive eigenvalues denoted by $\lambda_1,\cdots,\lambda_N>0$. Therefore, the eigenvalues of $\left(I+(WB)^{-1}\frac{\Sigma^{-1}}{\tau}\right)^{-1}$ are $\left(1+\frac{\lambda_1}{\tau}\right)^{-1},\cdots,\left(1+\frac{\lambda_N}{\tau}\right)^{-1}$, which are strictly less than one.

In summary, we have

$$\left\|\left(I+(WB)^{-1}\frac{\Sigma^{-1}}{\tau}\right)^{-1}B^{-1}W^{-1}QQ^{T}\right\|$$
$$\le\left\|\left(I+(WB)^{-1}\frac{\Sigma^{-1}}{\tau}\right)^{-1}\right\|\left\|B^{-1}W^{-1}QQ^{T}\right\|$$
$$\le\left(1+\frac{\lambda_N}{\tau}\right)^{-1}<1,$$

which suggests that the spectral norm of (S8) is bounded by $\left(1+\frac{\lambda_N}{\tau}\right)^{-1}$, from which we can see that a larger $\tau$ would generally drop the convergence rate. This is also confirmed by our simulation study (Figure S3), which shows that a higher-order approximation has much better performance (in terms of steps of convergence) for a larger $\tau$ (e.g., $\tau$>0.2). In addition, the spectral density of $\Sigma^{-1}$ also affects the convergence rate through $\lambda_N$. A large condition number of $\Sigma^{-1}$ would slow down the convergence, which is also corroborated by our simulation results (Figure S3). Higher-order approximation converged much faster under larger block sizes and stronger correlation, in which cases the condition number of the relatedness matrix is larger. This simulation also suggests that the first-order approximation is near-optimal for a common family-based design in which the average family size is 5 and most correlation coefficients are below 0.5.

### 3. Proof of the validity of the NR method for positive semidefinite covariance matrices

Here, we prove that using the GPPL $\tilde{l}_1$, the NR method is still valid for $\Sigma$ being SPSD except that the sum of elements in each row of $\Sigma$ is zero (i.e., $\Sigma$ has eigenvector $1$ with eigenvalue 0). We first show that when $\Sigma$ is SPSD and has eigenvector $1$ with eigenvalue 0, $V_{22}$, and thus $V$ become always non-invertible, which violates the regularity condition of the NR method. It is shown in A.2 that $WB-QQ^T$ is always positive semidefinite and has eigenvector $1$ with eigenvalue 0. Combined with the fact that $\Sigma^{-}1=0$ if $\Sigma 1=0$, we have

$$V_{22}\mathbf{1}=\left(\mathbf{WB}-\mathbf{QQ}^{\mathrm{T}}+\frac{\mathbf{\Sigma}^{-}}{\boldsymbol{\tau}}\right)\mathbf{1}$$
$$=\left(\mathbf{WB}-\mathbf{QQ}^{\mathrm{T}}\right)\mathbf{1}+\frac{\mathbf{\Sigma}^{-}}{\boldsymbol{\tau}}\mathbf{1}$$
$$=\mathbf{0},$$

which suggests that in such case, $V_{22}$ has a zero eigenvalue, and thus is non-invertible. Next, we show that $V_{22}$ is always invertible when $\Sigma$ is SPSD and $1$ is not one of its eigenvectors corresponding to the eigenvalue 0. We have shown in A.1 that

$$
\begin{aligned}
&WB - QQ^T \\
&= WB\left(I - B^{-1}\tilde{M}\tilde{A}^{2}\tilde{M}^{T}W\right) \\
&= WB(I - S_I S_{II})
\end{aligned},
$$

and $S_I S_{II}$ are product of two rectangular row stochastic matrices, which always has the largest eigenvalue one with eigenvector $\mathbf{1}$. Because $W$ and $\tilde{A}$ are diagonal matrices with only positive elements, and $\tilde{\mathbf{M}}$ is similar through permutation to a lower-triangular matrix when assuming no ties, it is easy to verify that the elements of $S_I S_{II}$ are all positive. Using Breslow's approximation for ties does not change this conclusion. According to the Perron–Frobenius theorem, $S_I S_{II}$ has a unique largest eigenvalue, which is one, which means that all the other vectors have its eigenvalue strictly less than one. Suppose that $\nu$ is an eigenvector of $V_{22}$. Consider its eigenvalue

$$
\begin{aligned}
&V_{22}\boldsymbol{\nu} \\
&= WB(I - S_I S_{II})\,\boldsymbol{\nu} + \frac{\boldsymbol{\Sigma}^{-1}}{\boldsymbol{\tau}}\boldsymbol{\nu}
\end{aligned}.
$$

If $\nu$ is $\mathbf{1}$, then the second term must be positive, and otherwise, the first term must be positive. Thus, the eigenvalues of $V_{22}$ are all positive, which suggests that the NR method is still valid.

### 4. Approximation of the log-determinant using the SLQ method

We describe the details of estimating the variance component $\tau$ when the relatedness matrix $\Sigma$ is large and fully dense. We estimate $\tau$ using the marginal likelihood

$$
l_2 = l_1(\hat{\boldsymbol{\theta}}) - \frac{1}{2}\log\left|\frac{J\left(l_1(\hat{\boldsymbol{\theta}}),\tau\right)}{2\pi}\right| = l_1(\hat{\boldsymbol{\theta}}) - \frac{1}{2}\log\left|\frac{\boldsymbol{W}(\hat{\boldsymbol{\theta}})\,\boldsymbol{B}(\hat{\boldsymbol{\theta}}) - \boldsymbol{Q}(\hat{\boldsymbol{\theta}})\,\boldsymbol{Q}(\hat{\boldsymbol{\theta}})^{T} + \frac{\boldsymbol{\Sigma}^{-1}}{\tau}}{2\pi}\right|, \quad \text{(S8)}
$$

where $\hat{\theta}$ is obtained by optimizing the PPL. Once $l_1$ is optimized, the addition step for estimating $\tau$ is the evaluation of the log-determinant in e.q.(S8), the time complexity of which is cubic when using a standard Cholesky decomposition. When $\Sigma^{-1}$ is dense and very large, this evaluation is computationally intensive. Therefore, we resort to a randomized method based on SLQ to reduce the time complexity to quadratic. We selected the SLQ method for approximating a log-determinant because our preliminary results suggested that it is more accurate than other randomized methods such as Chebyshev orthogonal polynomials (Han et al., 2016; Pace and LeSage, 2004), and Martin's Taylor expansion (Barry and Kelley Pace, 1999; Martin, 1992) under the same computational burden. The log-determinant to be approximated is

$$
\log\left|\boldsymbol{W}(\hat{\boldsymbol{\theta}})\,\boldsymbol{B}(\hat{\boldsymbol{\theta}}) - \boldsymbol{Q}(\hat{\boldsymbol{\theta}})\,\boldsymbol{Q}(\hat{\boldsymbol{\theta}})^{T} + \frac{\boldsymbol{\Sigma}^{-1}}{\tau}\right|. \quad \text{(S9)}
$$

The method works as follows. Given a certain matrix $\boldsymbol{P}$, we first approximate its log-determinant using a Monte Carlo trace estimator

$$\log|\boldsymbol{P}| = \mathrm{tr}(\log(\boldsymbol{P})) \approx \frac{1}{\mathrm{n_m}} \sum_{\mathrm{i}=1}^{\mathrm{n_m}} \mathbf{r}_{\mathrm{i}}^{\mathrm{T}} \log(\boldsymbol{P})\, \mathbf{r}_{\mathrm{i}},$$

where $\mathrm{n_m}$ is the number of Monte Carlo samples, and $r_i$ is an *i.i.d* sample from the Rademacher distribution as proposed in (Hutchinson, 1990). Since direct evaluation of $\log(\boldsymbol{P})$ is difficult, we rewrite it as

$$\begin{aligned}
&\frac{1}{\mathrm{n_m}} \sum_{\mathrm{i}=1}^{\mathrm{n_m}} \mathbf{r}_{\mathrm{i}}^{\mathrm{T}} \log(\boldsymbol{P})\, \mathbf{r}_{\mathrm{i}} \\
&= \frac{\mathbf{n}}{\mathbf{n_m}} \sum_{\mathrm{i}=1}^{\mathrm{n_m}} \frac{\mathbf{r_i}}{\sqrt{\mathbf{n}}}^{\mathrm{T}} \mathbf{U} \log(\Lambda)\, \mathbf{U}^{\mathrm{T}} \frac{\mathbf{r_i}}{\sqrt{\mathbf{n}}} \\
&= \frac{\mathbf{n}}{\mathbf{n_m}} \sum_{\mathrm{i}=1}^{\mathrm{n_m}} \tilde{\mathbf{r}}_{\mathrm{i}}^{\mathrm{T}} \log(\Lambda)\, \tilde{\mathbf{r}}_{\mathrm{i}} \\
&= \frac{\mathbf{n}}{\mathbf{n_m}} \sum_{\mathrm{i}=1}^{\mathrm{n_m}} \sum_{\mathrm{j}} \log\left(\lambda_{\mathrm{j}}\right) \tilde{\mathbf{r}}_{\mathrm{ij}}^{2}
\end{aligned},$$

where $\tilde{r}_i = U^T \frac{r_i}{\sqrt{n}}$, and $\tilde{r}_{ij}$ is the j[th] element in $\tilde{r}_i$. The second summation in the last line is recognized as a Riemann-Stieltjes integral, which can then be approximated using the Gauss quadrature rule

$$\sum_{\mathrm{j}} \log\left(\lambda_{\mathrm{j}}\right) \tilde{\mathrm{r}}_{\mathrm{ij}}^{2} = \int_{\lambda_{min}}^{\lambda_{max}} \log(\lambda)\, \mathrm{d}\tilde{\mathrm{r}}_{\mathrm{i}}^{2}(\lambda) \approx \sum_{k=1}^{n_q} \omega_{ik} \log\left(\varphi_{ik}\right),$$

where $n_q$ is the number of points in the Gauss quadrature rule, and $\omega_{ik}$ and $\varphi_{ik}$ are the weights and nodes in the Gauss quadrature rule to be determined. It is nicely shown in e.g., Theorem 4.1 in (Golub and Meurant, 2009), that the nodes $\varphi_{ik}$ are the zeros in the Lanczos orthogonal polynomial of $\boldsymbol{P}$, which are the eigenvalues of the tridiagonal matrix $T_{Lanczos}$ corresponding to the orthogonal polynomial (as shown in Theorem 6.2 in (Golub and Meurant, 2009)), which is the output of the Lanczos algorithm. The weights $\omega_{ik}$ are the square of the first element of the eigenvector $k$ of $T_{Lanczos}$. One potential issue is that the Lanczos algorithm is numerically unstable due to round-off errors. Even for a modest $n_q$, it is possible that the vectors produced by the algorithm become dependent. To prevent this issue, we stopped the algorithm once an off-diagonal element in $T_{Lanczos}$ was smaller than a small number (e.g., 1e-10).

The accuracy of the SLQ approximation highly depends on $n_q$ and $\mathrm{n_m}$. Although theoretical error bounds of the approximation of the log-determinant are given in (Ubaru et al., 2017), we investigated its empirical performance in COXMEG. Our results showed that $n_m = 100$ and $n_q = 10$ yielded highly accurate estimate of the variance component $\tau$ when the condition number of the relatedness matrix is not too large (Figure S6). We observed that the approximation was poor only in the scenario where the block size was large (500) and also the correlations were high (0.9) (Figure S6), which is relatively rarely encountered in a real data analysis.

#### 5. Simulation study for statistical power and empirical size

We investigated FPR and statistical power of the four methods, COXMEG-score, COXMEG-sparse, coxph with a shared frailty and coxme. Because dense matrices are overly time consuming for coxme, we investigated the power using block-diagonal covariance matrices, and expected that the conclusion should be generalized to dense covariance matrices. We first assessed the FPR under settings of different variance components, and correlation structure. We found that overall all methods except coxph controlled the type I error rate well. We found that COXMEG-score, COXMEG-sparse, and coxme had almost the same FPR as expected (Figure S1). In contrast, our results show that coxph with a shared frailty had inflated FPR when the correlation coefficients are 0.5, while the power is slightly diminished when the correlation coefficients are 0.1 (Figure S1). When the correlation coefficients are 0.9, coxph with a shared frailty controlled the FPR as well as the other methods (Figure S1), which makes sense because under this correlation, subjects within a block almost have the same random effect.

We next evaluated the statistical power for detecting the effect of a predictor using a simulation study. We considered multiple settings of different sample sizes, proportion of censoring, variance components, and correlation structure. More than 5000 subjects were needed to detect a log(HR) of 0.1. We observed that COXMEG-sparse shared almost the same statistical power as coxme, and COXMEG-score in all settings (Figure S2). We also noted that coxph with a shared frailty had very similar power compared to the other methods (Figure S2).

### B. Supplementary Tables

| CHR | POS | SNP | Gene | P-value |
|---|---|---|---|---|
| 1 | 161155392 | rs4575098 | ADAMTS4 | 9.420E-01 |
| 1 | 207786828 | rs6656401 | CR1 | 8.568E-03 |
| 2 | 127891427 | rs4663105 | BIN1 | 2.961E-04 |

| | | | | |
|---|---|---|---|---|
| 2 | 233981912 | rs10933431 | INPP5D | 1.154E-01 |
| 4 | 11026028 | rs6448453 | CLNK | 2.367E-02 |
| 6 | 32583357 | rs9269853 | HLA-DRB1 | 4.983E-02 |
| 6 | 47432637 | rs9381563 | CD2AP | 9.214E-01 |
| 7 | 99971834 | rs1859788 | ZCWPW1 | 3.128E-01 |
| 7 | 143108158 | rs11763230 | EPHA1 | 6.735E-01 |
| 8 | 27464929 | rs4236673 | CLU | 5.996E-02 |
| 10 | 11717397 | rs11257242 | ECHDC3 | 6.853E-01 |
| 11 | 59958380 | rs7935829 | MS4A6A | 3.141E-02 |
| 11 | 85776544 | rs10792832 | PICALM | 2.537E-03 |
| 14 | 92938855 | rs12590654 | SLC24A4 | 1.468E-01 |
| 15 | 59022615 | rs442495 | ADAM10 | 3.412E-01 |
| 15 | 63569902 | rs117618017 | APH1B | 6.624E-01 |
| 16 | 31133100 | rs59735493 | KAT8 | 6.414E-01 |
| 17 | 5138980 | rs113260531 | SCIMP | 7.153E-02 |

| | | | | |
|---|---|---|---|---|
| 17 | 47450775 | rs28394864 | ABI3 | 1.861E-02 |
| 19 | 1039323 | rs4147929 | ABCA7 | 3.925E-01 |
| 19 | 51727962 | rs3865444 | CD33 | 8.028E-01 |
| 20 | 54998544 | rs6014724 | CASS4 | 2.500E-02 |

Table S1. Results of 23 common SNPs identified by a recent meta-analysis of AD. The p-values of HRs were obtained from COXMEG-score based on a score test. The position is based on hg19.

### C. Supplementary Figures

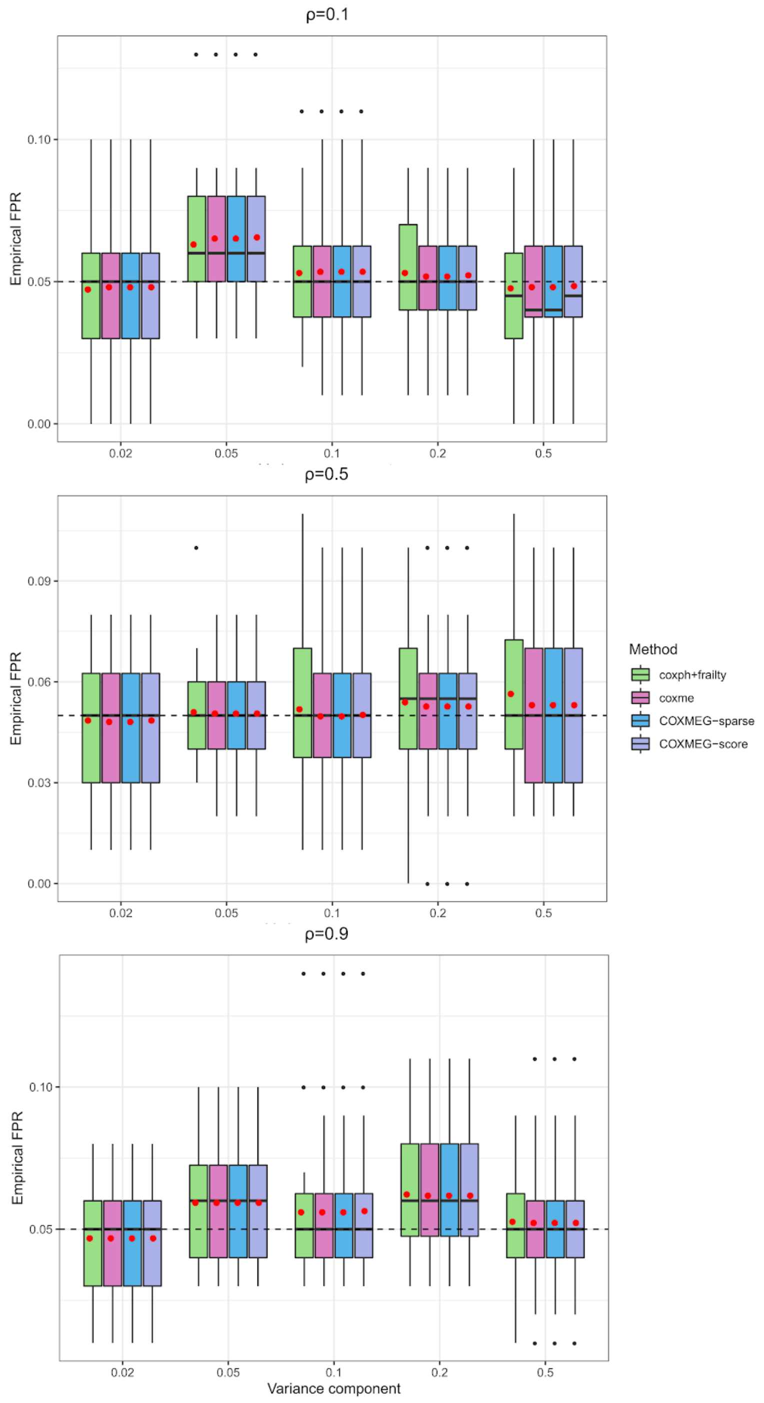


Figure S1. Comparison of empirical FPR of coxme, COXMEG-sparse, COXMEG-score, and coxph with a shared frailty. The relatedness matrix used in the simulation is a block-diagonal correlation matrix with the block size ranging between 5-100. We evaluated the FPR for the correlation ρ in each block being 0.1, 0.5, and 0.9, and the variance component $\tau$ between 0.02 and 0.5.

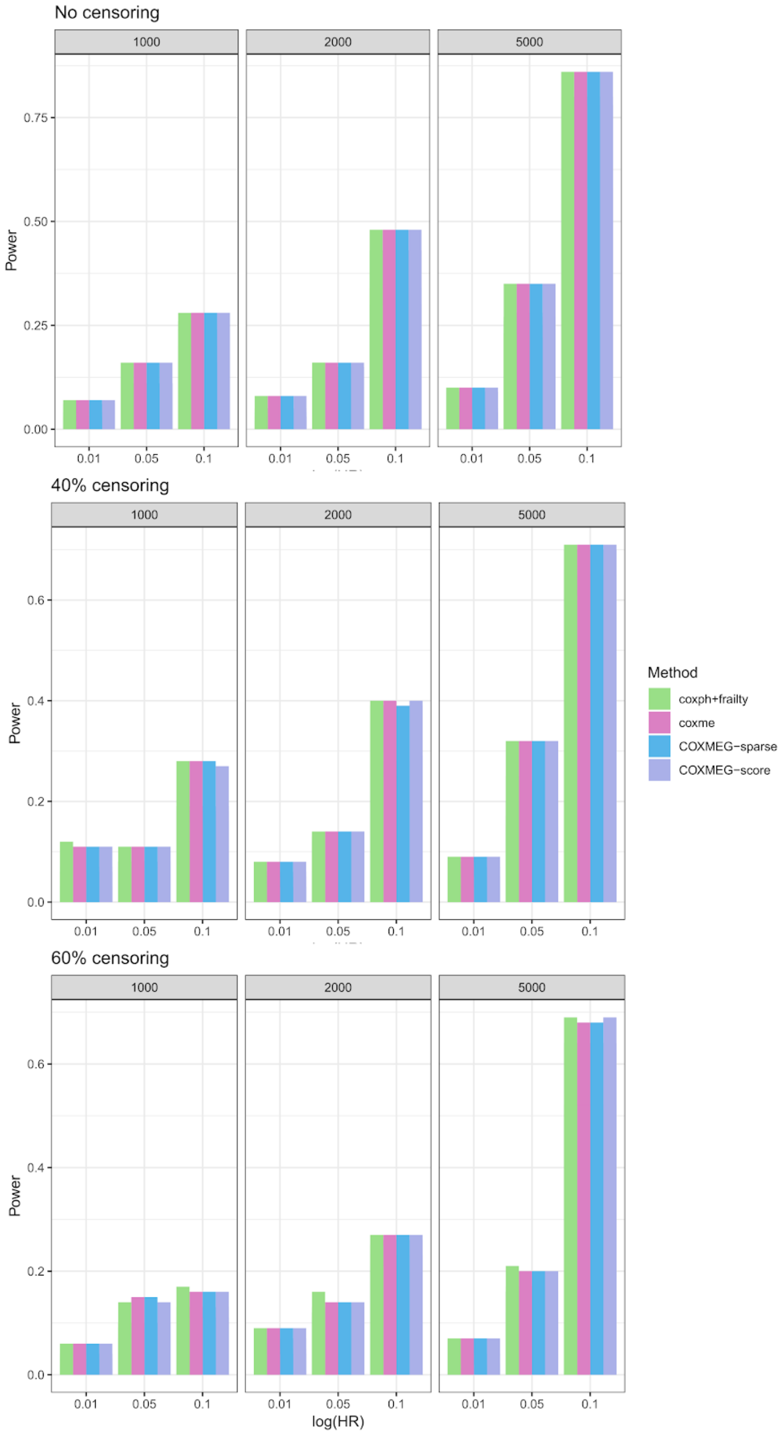


Figure S2. Comparison of empirical power of coxme, COXMEG-sparse, COXMEG-score, and coxph with a shared frailty. The relatedness matrix used in the simulation is a block-diagonal correlation matrix with the block size ranging between 5-100 and the correlation ρ ranging between 0.1 and 0.9. We evaluated the power for the HRs being 0.01, 0.05, and 0.1, and the sample size between 1000 and 5000. We also assess the power under no censoring, moderate censoring and heavy censoring.

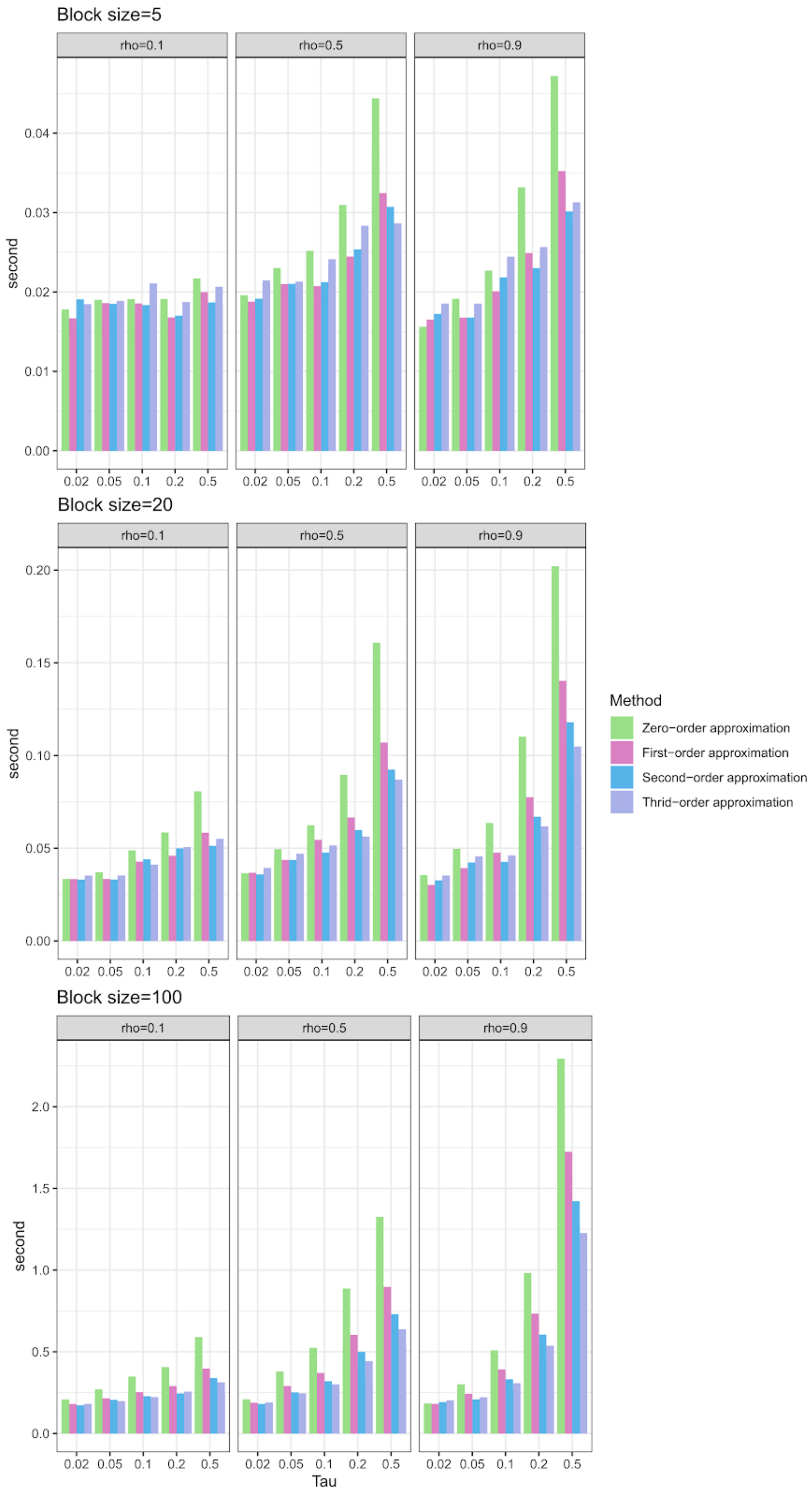


Figure S3. Evaluation of the convergence rate of higher-order approximations in COXMEG-sparse. The relatedness matrix used in the simulation is a block-diagonal correlation matrix with the block size ranging between 5-100 and the correlation ρ ranging between 0.1 and 0.9. For each setting, the convergence rate was measured by the time for estimating the HRs of one predictor given a variance component $\tau$ ranging from 0.02 to 0.5.

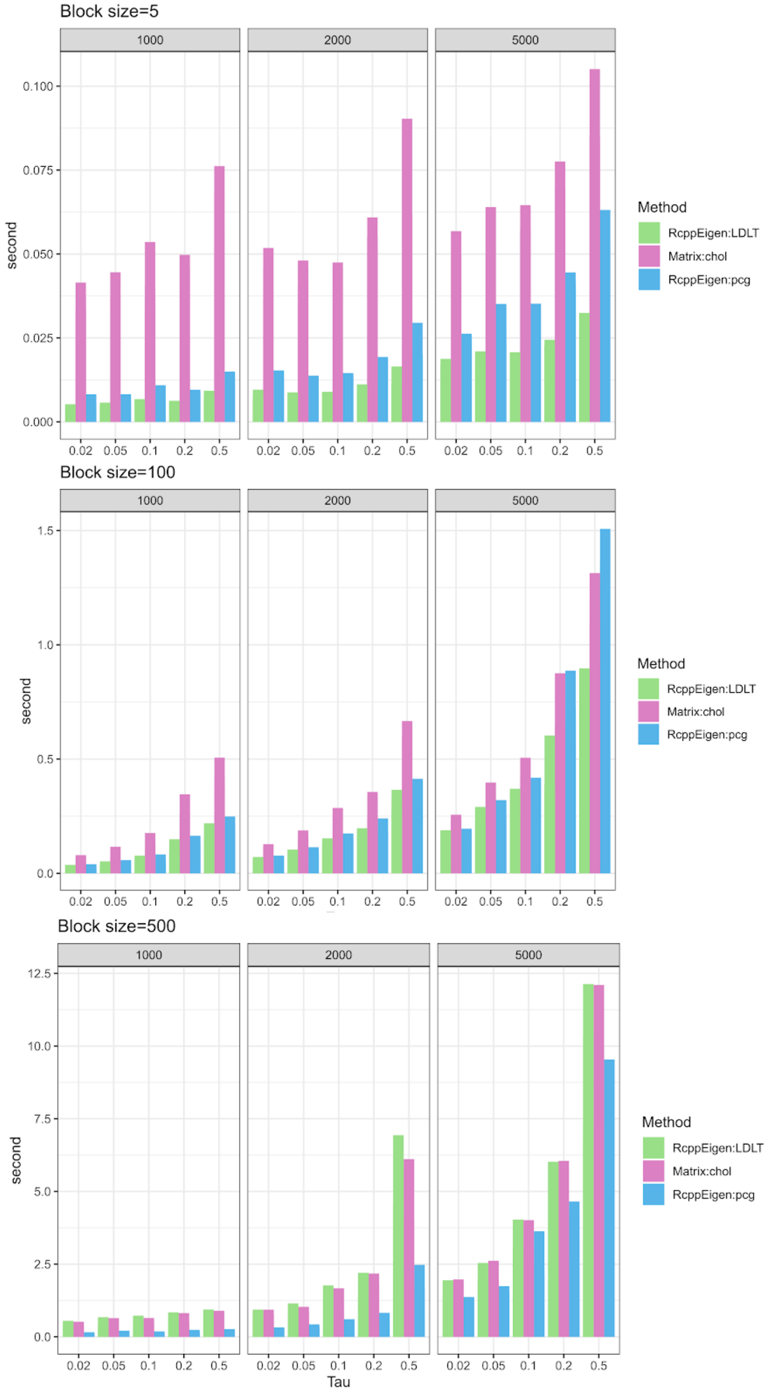


Figure S4. Evaluation of the computational performance of three methods (RcppEigen::LDLT, Matrix::solve using the Cholesky decomposition, and RcppEigen::CG with diagonal preconditioned) for solving the sparse linear system in COXMEG-sparse. The relatedness matrix used in the simulation is a block-diagonal correlation matrix with the block size varying between 5-500 and the correlation ρ being 0.5. For each setting, the convergence rate was measured by the time for estimating the HRs of one predictor given a variance component $\tau$ ranging from 0.02 to 0.5.

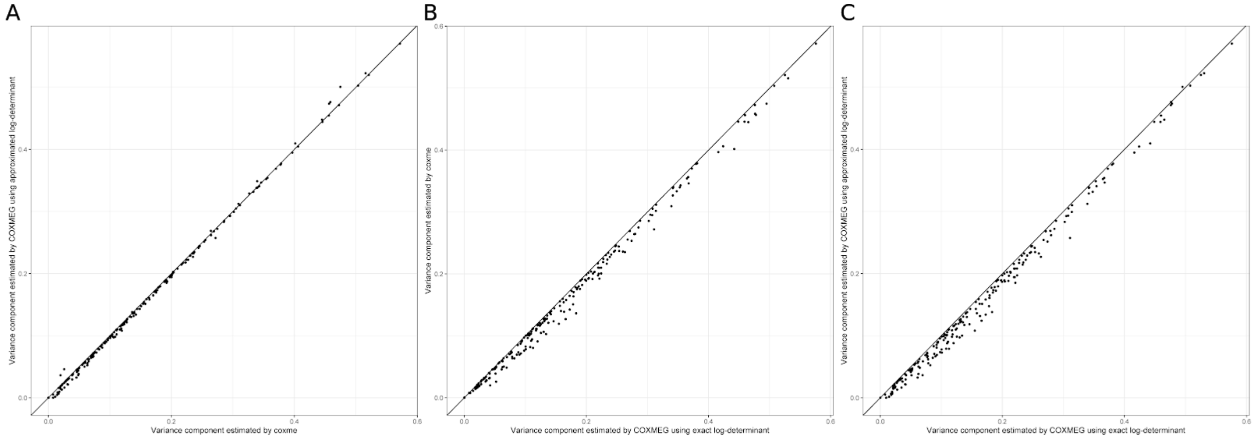


Figure S5. Comparison of estimated variance components by coxme, COXMEG-sparse with the exact log-determinant, and COXMEG-sparse with a diagonal approximated log-determinant. (A) COXMEG-sparse with a diagonal approximated log-determinant vs. coxme; (B) COXMEG-sparse with the exact log-determinant vs. coxme; (C) COXMEG-sparse with a diagonal approximated log-determinant vs. the exact log-determinant.

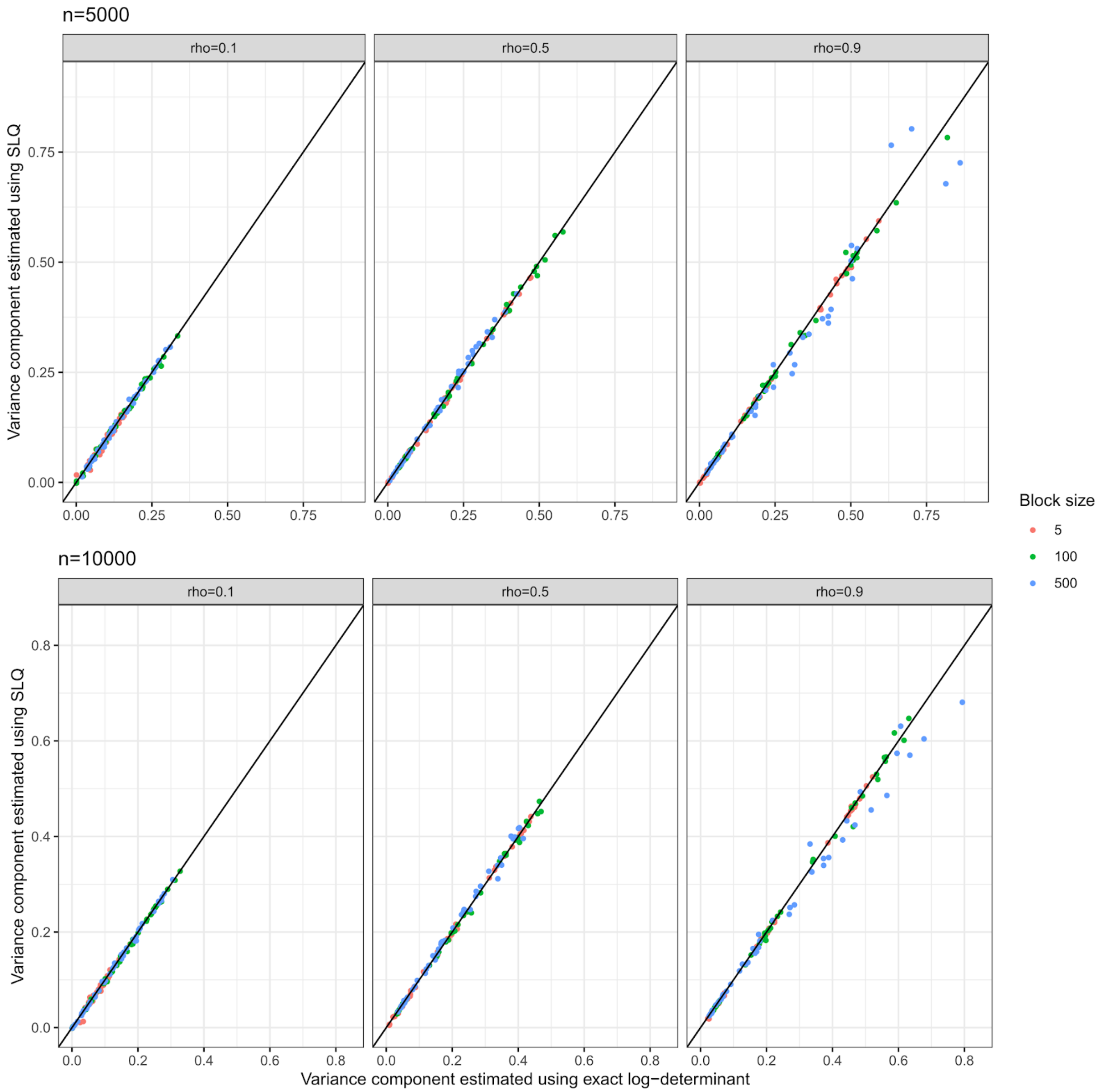


Figure S6. Comparison of estimated variance components by COXMEG-score with the exact log-determinant, and COXMEG-score with the SLQ approximation under sample sizes of 5000 and 10,000. The relatedness matrix used in the simulation is a block-diagonal correlation matrix with the block size varying between 5-500 and the correlation ρ between 0.1 and 0.9.

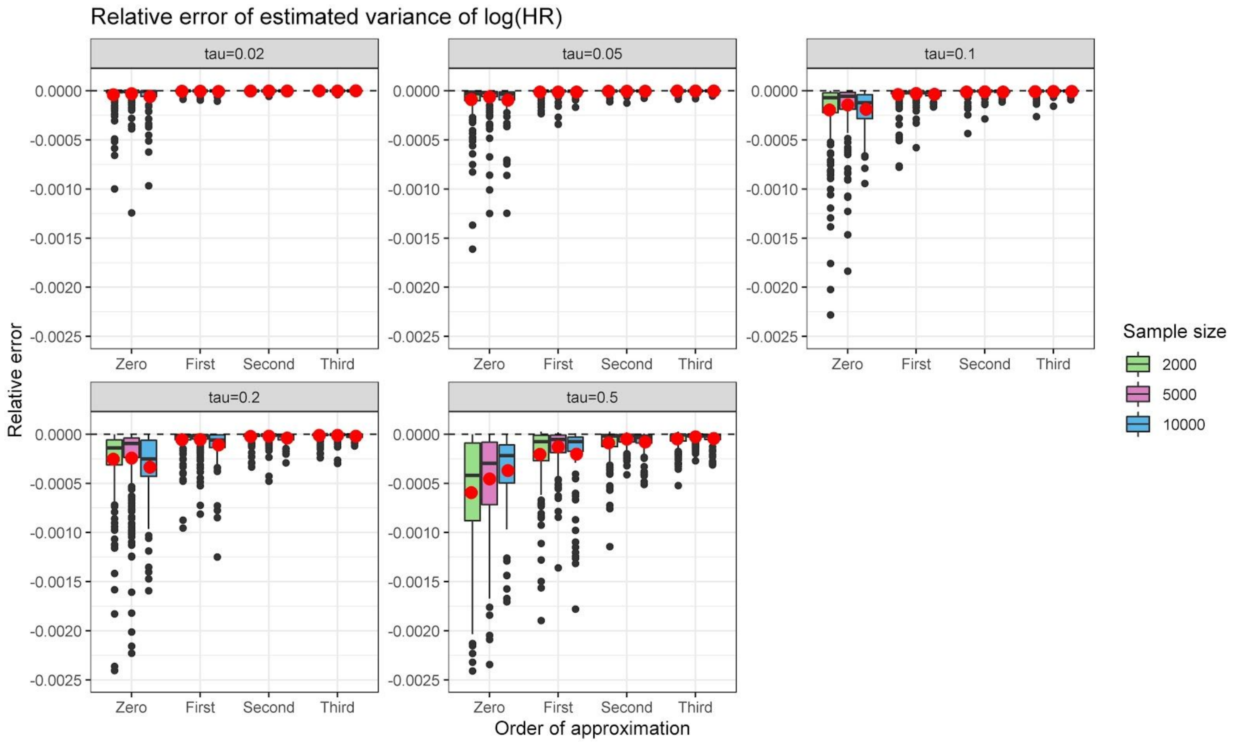


Figure S7. Relative error between the variance of log(HR) estimated using an approximated $V_{22}^{-1}$ in COXMEG-sparse and using an exact Hessian matrix. Four approximations of $V_{22}^{-1}$ (Zero-order to Third-order) were evaluated under settings of different sample sizes, and variance components. The red dots are the mean values.
