## Supplementary material for "Genome-wide association analysis of age-at-onset traits using Cox mixed-effects models": R_package_vignettes

*Liang He*

*2019-08-08*

### Overview

Time-to-event is one of the most important phenotypes in genetic epidemiology. The R-package, “coxmeg”, provides a set of utilities to fit a Cox mixed-effects model and to efficiently perform genome-wide association analysis of time-to-event phenotypes using a Cox mixed-effects model.

### Installation

#### Most-recent version

```
install.packages("coxmeg", repos="http://R-Forge.R-project.org")
```

### Fit a Cox mixed-effects model with a sparse relatedness matrix

We illustrate how to use coxmeg to fit a Cox mixed-effects model with a sparse relatedness matrix. We first simulate a block-diagonal relatedness matrix for a cohort consisting of 200 families, each of which has five members.

```
library(coxmeg)
```

```
## Loading required package: Rcpp
```

```
library(MASS)
library(Matrix)
n_f <- 200
mat_list <- list()
size <- rep(5,n_f)
offd <- 0.5
for(i in 1:n_f)
{
  mat_list[[i]] <- matrix(offd,size[i],size[i])
  diag(mat_list[[i]]) <- 1
}
```

```
sigma <- as.matrix(bdiag(mat_list))
sigma = as(sigma,'dgCMatrix')
```

We use ‘dgCMatrix’ to save memory. Next, we simulate random effects and time-to-event outcomes assuming a constant baseline hazard function. We assume that the variance component is 0.2. We also simulate a risk factor with log(HR)=0.1.

```
n = nrow(sigma)
tau_var <- 0.2
x <- mvrnorm(1, rep(0,n), tau_var*sigma)
pred = rnorm(n,0,1)
myrates <- exp(x+0.1*pred-1)
y <- rexp(n, rate = myrates)
cen <- rexp(n, rate = 0.02 )
ycen <- pmin(y, cen)
outcome <- cbind(ycen,as.numeric(y <= cen))
head(outcome)
```

```
##              ycen
## [1,]  1.1965156 1
## [2,]  0.7331514 1
## [3,]  0.3306775 1
## [4,]  0.3456029 1
## [5,]  2.9256348 1
## [6,] 10.2343296 1
```

```
sigma[1:5,1:5]
```

```
## 5 x 5 sparse Matrix of class "dgCMatrix"
##
## [1,] 1.0 0.5 0.5 0.5 0.5
## [2,] 0.5 1.0 0.5 0.5 0.5
## [3,] 0.5 0.5 1.0 0.5 0.5
## [4,] 0.5 0.5 0.5 1.0 0.5
## [5,] 0.5 0.5 0.5 0.5 1.0
```

We fit a Cox mixed-effects model using the `coxmeg` function. We set `dense=FALSE` to indicate that the relatedness matrix is sparse. However, the function will automatically treat it as dense if there are more than 50% non-zero elements in the matrix. We set `order=1` to use the first-order approximation of the inverse Hessian matrix in the optimization.

```
re = coxmeg(outcome,sigma,pred,order=1,dense=FALSE)
```

```
## [1] "Remove 0 subjects censored before the first failure."
## [1] "The sample size included is 1000."
## [1] "The relatedness matrix is treated as sparse."
```

```
re
```

```
## $beta
##             [,1]
## [1,] 0.07710682
##
## $HR
##           [,1]
## [1,] 1.080157
##
## $sd_beta
## [1] 0.03770179
##
## $p
##             [,1]
## [1,] 0.04083743
##
## $tau
## [1] 0.2765141
##
## $iter
## [1] 16
##
## $rank
## [1] 1000
##
## $nsam
## [1] 1000
##
## $int_ll
## [1] 11491
```

In the above result, `tau` is the estimated variance component, and `int_ll` is -2*log(lik) of the integrated/marginal likelihood of tau.

```
library(coxme)
```

```
## Warning: package 'coxme' was built under R version 3.5.3
```

```
## Loading required package: survival
```

```
## Loading required package: bdsmatrix
```

```
##
## Attaching package: 'bdsmatrix'
```

```
## The following object is masked from 'package:base':
##
##     backsolve
```

```
bls <- c(1)
for(i in (size[1]-1):1)
{bls <- c(bls, c(rep(offd,i),1))}
tmat <- bdsmatrix(blocksize=size,
blocks=rep(bls,n_f),dimnames=list(as.character(1:n),as.character(1:n)))
re_coxme = coxme(Surv(outcome[,1],outcome[,2])~as.matrix(pred)+(1|as.character(1:n)),
varlist=list(tmat),ties='breslow')
re_coxme
```

```
## Cox mixed-effects model fit by maximum likelihood
##
##   events, n = 945, 1000
##   Iterations= 7 35
##                     NULL  Integrated    Fitted
## Log-likelihood -5588.539  -5577.983 -5392.659
##
##                    Chisq     df          p    AIC     BIC
## Integrated loglik  21.11   2.00 2.6051e-05 17.11    7.41
##  Penalized loglik 391.76 166.09 0.0000e+00 59.58 -746.14
##
## Model:  Surv(outcome[, 1], outcome[, 2]) ~ as.matrix(pred) + (1 | as.character(1:n))
## Fixed coefficients
##                       coef exp(coef)   se(coef)    z     p
## as.matrix(pred) 0.07719058  1.080248 0.03773991 2.05 0.041
##
## Random effects
##  Group              Variable Std Dev   Variance
##  as.character.1.n. Vmat.1   0.5277798 0.2785515
```

We compare the results with coxme. The integrated log-likelihoods cannot be compared directly because different approximation of log-determinant is used.

### Perform GWAS of an age-at-onset phenotype with a sparse relatedness matrix

We illustrate how to perform a GWAS using the `coxmeg_plink` function. This function supports plink bed files. We provide example files in the package. The example plink files include 20 SNPs and 3000 subjects from 600 families. The following code performs a GWAS for all SNPs in the example bed files.

```
library(coxmeg)
bed = system.file("extdata", "example_null.bed", package = "coxmeg")
bed = substr(bed,1,nchar(bed)-4)
pheno = system.file("extdata", "ex_pheno.txt", package = "coxmeg")
cov = system.file("extdata", "ex_cov.txt", package = "coxmeg")

## building a relatedness matrix
n_f <- 600
mat_list <- list()
size <- rep(5,n_f)
offd <- 0.5
```

```
for(i in 1:n_f)
{
  mat_list[[i]] <- matrix(offd,size[i],size[i])
  diag(mat_list[[i]]) <- 1
}
sigma <- as.matrix(bdiag(mat_list))

re = coxmeg_plink(pheno,sigma,bed,cov=cov,detap=TRUE,dense=FALSE,verbose=FALSE)
```

```
## Excluding 0 SNP on non-autosomes
## Excluding 0 SNP (monomorphic: TRUE, MAF: 0.05, missing rate: 0)
```

```
re
```

```
## $summary
##     snp.id chromosome position allele      afreq   index         beta
## 1   null_0          1        1    d/D 0.30983333  null_0  0.015672101
## 2   null_1          1        2    d/D 0.23466667  null_1  0.019439150
## 3   null_2          1        3    D/d 0.14033333  null_2 -0.049845757
## 4   null_3          1        4    D/d 0.16183333  null_3  0.044130767
## 5   null_4          1        5    d/D 0.19933333  null_4  0.028473176
## 6   null_5          1        6    D/d 0.11800000  null_5 -0.114319159
## 7   null_6          1        7    d/D 0.09483333  null_6 -0.017981231
## 8   null_7          1        8    D/d 0.49683333  null_7 -0.004207897
## 9   null_8          1        9    d/D 0.31366667  null_8 -0.063741849
## 10  null_9          1       10    D/d 0.49183333  null_9 -0.008409562
## 11 null_10          1       11    d/D 0.34833333 null_10 -0.013581479
## 12 null_11          1       12    D/d 0.25100000 null_11  0.037508301
## 13 null_12          1       13    d/D 0.17500000 null_12 -0.017215848
## 14 null_13          1       14    D/d 0.06333333 null_13 -0.068207724
## 15 null_14          1       15    D/d 0.20833333 null_14 -0.013965386
## 16 null_15          1       16    d/D 0.17050000 null_15  0.002172773
## 17 null_16          1       17    D/d 0.33550000 null_16  0.004762350
## 18 null_17          1       18    d/D 0.26633333 null_17  0.001786995
## 19 null_18          1       19    D/d 0.09433333 null_18 -0.016052310
## 20 null_19          1       20    d/D 0.11650000 null_19 -0.022398126
##           HR    sd_beta           p
## 1  1.0157956 0.02938524 0.593803537
## 2  1.0196293 0.03222054 0.546298835
## 3  0.9513762 0.03860368 0.196628160
## 4  1.0451190 0.03701019 0.233106387
## 5  1.0288824 0.03432500 0.406811816
## 6  0.8919732 0.04234095 0.006934636
## 7  0.9821795 0.04655562 0.699325464
## 8  0.9958009 0.02717805 0.876957699
## 9  0.9382472 0.02958441 0.031195036
## 10 0.9916257 0.02730686 0.758108827
## 11 0.9865103 0.02859980 0.634872392
## 12 1.0382206 0.03113254 0.228282858
## 13 0.9829315 0.03628637 0.635183349
## 14 0.9340664 0.05698849 0.231357835
```

```
## 15 0.9861317 0.03431600 0.684034201
## 16 1.0021751 0.03685682 0.952990554
## 17 1.0047737 0.02859957 0.867749134
## 18 1.0017886 0.03098518 0.954009439
## 19 0.9840758 0.04731969 0.734435643
## 20 0.9778508 0.04231689 0.596600710
##
## $tau
## [1] 0.04028041
##
## $rank
## [1] 3000
##
## $nsam
## [1] 3000
```

The above code first retrieves the full path of the files. If the full path is not given, the function will search the current working directory. The file name for the bed file should not include the suffix (.bed). The phenotype and covariate files have the same format as used in plink, and the IDs must be consistent with the bed files. Specifically, the phenotype file should include four columns including family ID, individual ID, time, and status. The covariate file always starts with two columns, family ID and individual ID. Missing values in the phenotype and covariate files are denoted by -9 and NA, respectively. In the current version, the `coxmeg_plink` function does not impute genotypes itself, and only SNPs without missing values will be analyzed, so it will be better to use imputed genotype data.

The `coxmeg_plink` function fist estimates the variance component with only the covariates, and then uses it to analyze each SNP after filtering. These two steps can be done separately as follows. The first command without bed only esitmates the variance component tau, and the second command uses the estimated tau to analyze the SNPs. The `coxmeg_plink` function will write a temporary .gds file for the SNPs in the same folder of the bed files. The temporary file is removed after the analysis is done.

```
re = coxmeg_plink(pheno,sigma,cov=cov,detap=TRUE,dense=FALSE,verbose=FALSE)
re
```

```
## $tau
## [1] 0.04028041
##
## $iter
## [1] 15
##
## $rank
## [1] 3000
##
## $nsam
## [1] 3000
```

```
re = coxmeg_plink(pheno,sigma,bed,tau=re$tau,cov=cov,detap=TRUE,dense=FALSE,verbose=FALSE)
```

```
## Excluding 0 SNP on non-autosomes
## Excluding 0 SNP (monomorphic: TRUE, MAF: 0.05, missing rate: 0)
```

```
re
```

```
## $summary
##     snp.id chromosome position allele      afreq   index         beta
## 1   null_0          1        1    d/D 0.30983333  null_0  0.015672101
## 2   null_1          1        2    d/D 0.23466667  null_1  0.019439150
## 3   null_2          1        3    D/d 0.14033333  null_2 -0.049845757
## 4   null_3          1        4    D/d 0.16183333  null_3  0.044130767
## 5   null_4          1        5    d/D 0.19933333  null_4  0.028473176
## 6   null_5          1        6    D/d 0.11800000  null_5 -0.114319159
## 7   null_6          1        7    d/D 0.09483333  null_6 -0.017981231
## 8   null_7          1        8    D/d 0.49683333  null_7 -0.004207897
## 9   null_8          1        9    d/D 0.31366667  null_8 -0.063741849
## 10  null_9          1       10    D/d 0.49183333  null_9 -0.008409562
## 11 null_10          1       11    d/D 0.34833333 null_10 -0.013581479
## 12 null_11          1       12    D/d 0.25100000 null_11  0.037508301
## 13 null_12          1       13    d/D 0.17500000 null_12 -0.017215848
## 14 null_13          1       14    D/d 0.06333333 null_13 -0.068207724
## 15 null_14          1       15    D/d 0.20833333 null_14 -0.013965386
## 16 null_15          1       16    d/D 0.17050000 null_15  0.002172773
## 17 null_16          1       17    D/d 0.33550000 null_16  0.004762350
## 18 null_17          1       18    d/D 0.26633333 null_17  0.001786995
## 19 null_18          1       19    D/d 0.09433333 null_18 -0.016052310
## 20 null_19          1       20    d/D 0.11650000 null_19 -0.022398126
##           HR    sd_beta           p
## 1  1.0157956 0.02938524 0.593803537
## 2  1.0196293 0.03222054 0.546298835
## 3  0.9513762 0.03860368 0.196628160
## 4  1.0451190 0.03701019 0.233106387
## 5  1.0288824 0.03432500 0.406811816
## 6  0.8919732 0.04234095 0.006934636
## 7  0.9821795 0.04655562 0.699325464
## 8  0.9958009 0.02717805 0.876957699
## 9  0.9382472 0.02958441 0.031195036
## 10 0.9916257 0.02730686 0.758108827
## 11 0.9865103 0.02859980 0.634872392
## 12 1.0382206 0.03113254 0.228282858
## 13 0.9829315 0.03628637 0.635183349
## 14 0.9340664 0.05698849 0.231357835
## 15 0.9861317 0.03431600 0.684034201
## 16 1.0021751 0.03685682 0.952990554
## 17 1.0047737 0.02859957 0.867749134
## 18 1.0017886 0.03098518 0.954009439
## 19 0.9840758 0.04731969 0.734435643
## 20 0.9778508 0.04231689 0.596600710
## 
## $tau
## [1] 0.04028041
## 
## $rank
## [1] 3000
## 
```

```
## $nsam
## [1] 3000
```

### Handle positive semidefinite relatedness matrices

We now assume that the first two subjects in the sample are monozygotic twins, and thus the relatedness matrix becomes positive semidefinite. To handle this, use `spd=FALSE` as follows.

```
sigma[2,1] = sigma[1,2] = 1
re = coxmeg_plink(pheno,sigma,cov=cov,detap=TRUE,dense=FALSE,verbose=FALSE,spd=FALSE)
```

```
## Warning in chol.default(x, pivot = TRUE): the matrix is either rank-
## deficient or indefinite
```

```
re
```

```
## $tau
## [1] 0.04038302
##
## $iter
## [1] 15
##
## $rank
## [1] 2999
##
## $nsam
## [1] 3000
```

The warning indicates that the matrix is not full rank. The rank is less than the sample size.

### Perform GWAS of an age-at-onset phenotype with a dense relatedness matrix

When the relatedness matrix is dense, it will be more efficient to specify `dense=TRUE`, and use preconditioned conjugate gradiant `solver=2` and stochastic lanczos quadrature `detap=TRUE` in the optimization. These can be specified as follows.

```
re = coxmeg_plink(pheno,sigma,bed,cov=cov,detap=TRUE,dense=TRUE,verbose=FALSE,spd=FALSE,solver=2)
```

```
## Warning in chol.default(x, pivot = TRUE): the matrix is either rank-
## deficient or indefinite
```

```
## Excluding 0 SNP on non-autosomes
## Excluding 0 SNP (monomorphic: TRUE, MAF: 0.05, missing rate: 0)
```

```
re
```

```
## $summary
##     snp.id chromosome position allele      afreq   index         beta
## 1   null_0          1        1    d/D 0.30983333  null_0  0.015838551
## 2   null_1          1        2    d/D 0.23466667  null_1  0.019181203
## 3   null_2          1        3    D/d 0.14033333  null_2 -0.050919545
## 4   null_3          1        4    D/d 0.16183333  null_3  0.043805519
## 5   null_4          1        5    d/D 0.19933333  null_4  0.028520372
## 6   null_5          1        6    D/d 0.11800000  null_5 -0.114786460
## 7   null_6          1        7    d/D 0.09483333  null_6 -0.017753886
## 8   null_7          1        8    D/d 0.49683333  null_7 -0.004390952
## 9   null_8          1        9    d/D 0.31366667  null_8 -0.063735344
## 10  null_9          1       10    D/d 0.49183333  null_9 -0.008631232
## 11 null_10          1       11    d/D 0.34833333 null_10 -0.013572384
## 12 null_11          1       12    D/d 0.25100000 null_11  0.037784616
## 13 null_12          1       13    d/D 0.17500000 null_12 -0.017103035
## 14 null_13          1       14    D/d 0.06333333 null_13 -0.068366813
## 15 null_14          1       15    D/d 0.20833333 null_14 -0.014516846
## 16 null_15          1       16    d/D 0.17050000 null_15  0.001953720
## 17 null_16          1       17    D/d 0.33550000 null_16  0.004532554
## 18 null_17          1       18    d/D 0.26633333 null_17  0.001434312
## 19 null_18          1       19    D/d 0.09433333 null_18 -0.016109412
## 20 null_19          1       20    d/D 0.11650000 null_19 -0.023249652
##           HR    sd_beta           p
## 1  1.0159646 0.02947788 0.591058421
## 2  1.0193663 0.03233051 0.552990481
## 3  0.9503551 0.03871264 0.188402093
## 4  1.0447791 0.03714376 0.238258512
## 5  1.0289310 0.03444224 0.407634338
## 6  0.8915565 0.04251088 0.006930472
## 7  0.9824028 0.04670473 0.703848670
## 8  0.9956187 0.02728894 0.872167521
## 9  0.9382533 0.02965122 0.031594645
## 10 0.9914059 0.02739107 0.752677283
## 11 0.9865193 0.02868787 0.636138073
## 12 1.0385075 0.03122816 0.226296793
## 13 0.9830424 0.03640299 0.638480052
## 14 0.9339178 0.05715499 0.231632037
## 15 0.9855880 0.03444428 0.673420033
## 16 1.0019556 0.03697487 0.957860085
## 17 1.0045428 0.02870876 0.874550961
## 18 1.0014353 0.03110110 0.963216401
## 19 0.9840197 0.04743620 0.734156528
## 20 0.9770185 0.04247942 0.584161879
##
## $tau
## [1] 0.04454584
##
## $rank
## [1] 2999
##
## $nsam
```

```
## [1] 3000
```

The above command estimates HRs and report p-values. Instead, a score test, which is computationally much more efficient, can also be used by specifying `score=TRUE`.

```
re =
coxmeg_plink(pheno,sigma,bed,tau=re$tau,cov=cov,detap=TRUE,dense=TRUE,verbose=FALSE,spd=FALSE,solver=2,
)
```

```
## Warning in chol.default(x, pivot = TRUE): the matrix is either rank-
## deficient or indefinite
```

```
## Excluding 0 SNP on non-autosomes
## Excluding 0 SNP (monomorphic: TRUE, MAF: 0.05, missing rate: 0)
```

```
re
```

```
## $summary
##     snp.id chromosome position allele      afreq   index  score_test
## 1    null_0          1        1    d/D 0.30983333  null_0 0.288760652
## 2    null_1          1        2    d/D 0.23466667  null_1 0.351994465
## 3    null_2          1        3    D/d 0.14033333  null_2 1.728992384
## 4    null_3          1        4    D/d 0.16183333  null_3 1.391405959
## 5    null_4          1        5    d/D 0.19933333  null_4 0.686075221
## 6    null_5          1        6    D/d 0.11800000  null_5 7.310979969
## 7    null_6          1        7    d/D 0.09483333  null_6 0.144577647
## 8    null_7          1        8    D/d 0.49683333  null_7 0.025925708
## 9    null_8          1        9    d/D 0.31366667  null_8 4.614335982
## 10   null_9          1       10    D/d 0.49183333  null_9 0.099339183
## 11 null_10          1       11    d/D 0.34833333 null_10 0.223817206
## 12 null_11          1       12    D/d 0.25100000 null_11 1.464121599
## 13 null_12          1       13    d/D 0.17500000 null_12 0.220788370
## 14 null_13          1       14    D/d 0.06333333 null_13 1.431068880
## 15 null_14          1       15    D/d 0.20833333 null_14 0.177643651
## 16 null_15          1       16    d/D 0.17050000 null_15 0.002794204
## 17 null_16          1       17    D/d 0.33550000 null_16 0.024954283
## 18 null_17          1       18    d/D 0.26633333 null_17 0.002127861
## 19 null_18          1       19    D/d 0.09433333 null_18 0.115193735
## 20 null_19          1       20    d/D 0.11650000 null_19 0.299800342
##              p
## 1  0.591015831
## 2  0.552986277
## 3  0.188539631
## 4  0.238167810
## 5  0.407502560
## 6  0.006853454
## 7  0.703771964
## 8  0.872081886
## 9  0.031705768
```

```
## 10 0.752624086
## 11 0.636146665
## 12 0.226275468
## 13 0.638439875
## 14 0.231590076
## 15 0.673406119
## 16 0.957843273
## 17 0.874481032
## 18 0.963207635
## 19 0.734306892
## 20 0.584007615
##
## $tau
## [1] 0.04454584
##
## $rank
## [1] 2999
##
## $nsam
## [1] 3000
```
